# A comparison of neuronal population dynamics measured with calcium imaging and electrophysiology

**DOI:** 10.1101/840686

**Authors:** Ziqiang Wei, Bei-Jung Lin, Tsai-Wen Chen, Kayvon Daie, Karel Svoboda, Shaul Druckmann

## Abstract

Calcium imaging with fluorescent protein sensors is widely used to record activity in neuronal populations. The transform between neural activity and calcium-related fluorescence involves nonlinearities and a low-pass filter, but the effects of the transformation on analyses of neural populations are not well understood. We compared neuronal spikes and fluorescence in matched neural populations in behaving mice. We report multiple discrepancies between analyses performed on the two types of data, which were only partially resolved by spike inference algorithms applied to fluorescence. To model the relation between spiking and fluorescence we simultaneously recorded spikes and fluorescence from individual neurons. Using these recordings we developed a model transforming spike trains to synthetic-imaging data. The model recapitulated the differences in analyses. Our analysis highlights challenges in relating electrophysiology and imaging data, and suggests forward modeling as an effective way to understand differences between these data.

## Introduction

Electrophysiological recordings (‘ephys’) and calcium imaging offer distinct tradeoffs for interrogating activity in neural populations. Ephys directly reports spiking of neurons with high signal-to-noise ratio, temporal fidelity, and dynamic range, but typically offers access only to a sparse subset of relatively active neurons (Buzsaki, 2004). In addition, the ability to track the same population of neurons across time, important for understanding the neural basis of learning, remains challenging (Dhawale et al., 2017; Ganguly and Carmena, 2009; Tolias et al., 2007).

Calcium imaging provides access to large numbers of neurons simultaneously (Huber et al., 2012; Peron et al., 2015b; Sofroniew et al., 2016; Stringer et al., 2019), potentially with cell type specificity (Fu et al., 2014; Peron et al., 2015a). Moreover, calcium imaging can track the activity of the same neuronal populations over time (Huber et al., 2012; Peters et al., 2014). Indeed, with the development of sensitive fluorescent protein-based indicators (Akerboom et al., 2012; Chen et al., 2013; Dana et al., 2016; Dana et al., 2018; Inoue et al., 2015; Ohkura et al., 2012; Tian et al., 2009) and powerful new imaging methods (Hamel et al., 2015; Sofroniew et al., 2016) calcium imaging has been rapidly adopted for measurements of neural population activity.

However, calcium imaging reports spikes only indirectly (Grienberger and Konnerth, 2012; Peron et al., 2015a). The transformation from spikes to calcium is inherently non-linear due to the dynamics of intracellular calcium concentrations (Scheuss et al., 2006). Additional nonlinearities are imposed by the protein-based indicators of calcium (Akerboom et al., 2012; Chen et al., 2013; Pologruto et al., 2004; Tian et al., 2009). Together these produce a low-pass filtered, delayed, and transformed version of neural activity, which complicates relating neural activity to behavior. Calcium imaging also has lower signal-to-noise ratio for detecting spikes and limited dynamic range (Peron et al., 2015a). In addition, during animal behavior, spike rates can vary by orders of magnitude across behavioral epochs and across neurons, even neurons of the same type (Hromadka et al., 2008; Li et al., 2015; O’Connor et al., 2010) and spike rates change over times of milliseconds to seconds (Brody et al., 2003; Li et al., 2015; Li et al., 2016). Finally, the coupling between spikes and calcium-dependent fluorescence likely differs across different neuron types and even individual neurons within a type (Chen et al., 2013; Maravall et al., 2000).

The complexities in the relation between spiking activity and calcium imaging at the level of single neurons have been long appreciated (Akerboom et al., 2012; Chen et al., 2013; Greenberg et al., 2018; Pologruto et al., 2004; Scheuss et al., 2006; Tian et al., 2009). However, the effect of these factors on analyses of population activity are not yet fully known (Cunningham and Yu, 2014). Ideally a detailed understanding of the transformation from spikes-to-calcium-dependent fluorescence would allow inversion of this transformation and the reliable extraction of spikes. Calcium indicators with high sensitivity allow reliable detection of action potentials, at least under conditions when single spikes or burst of spikes are separated in time (Chen et al., 2013; Dana et al., 2014; Theis et al., 2016). However, under behaviorally relevant conditions neurons operate with a large range of spike rates, and spiking responses are typically superposed on a substantial background spike rate, which varies across the population (Li et al., 2015; O’Connor et al., 2010). Moreover, neuron-to-neuron variability in calcium dynamics, calcium indicator dynamics and patterns of firing rate could conspire to make this inversion challenging. These issues are compounded by the paucity of simultaneously recorded spikes and fluorescence data. It is therefore unclear if spike inference can invert the fluorescence data accurately to eliminate potential discrepancies between analyses performed on ephys and calcium imaging data.

Here, we explore these issues empirically in the context of a decision-making task, where the dynamics of the neural circuit are rich and variable across neurons. In particular, neurons in frontal cortex show a wide range of spike rates and exhibit diverse temporal dynamics and selectivity (Brody et al., 2003; Li et al., 2015; Wei et al., 2019). We analyzed ephys and calcium imaging measured in matched neuronal populations in the same delayed response task and directly compared the results of standard measurements of selectivity and population dynamics. We find qualitative discrepancies at both the level of single cells and neural populations. Spike inference algorithms were limited in resolving these differences. However, a phenomenological model of the spike-to-fluorescence transformation, based on simultaneous imaging and electrophysiology data (Akerboom et al., 2012; Chen et al., 2013), explains many differences across the data sets. Finally, we developed a web-based platform, im-phys.org, that allows quantification of the effects of various transformations from electrophysiology to imaging.

## Results

We measured neural activity using electrophysiology (‘ephys’) and calcium imaging under identical behavioral conditions and in matched neural populations, but in separate experiments. Mice performed a tactile delayed response task (Guo et al., 2014a; Guo et al., 2014b; Li et al., 2015) (Figure 1A). In each trial, mice judged the location of an object with their whiskers. During the subsequent delay epoch (approximately 1.3 seconds), mice planned an upcoming response. Following an auditory ‘go’ cue, mice reported object location with directional licking (lick-left or lick-right).

**Figure 1.**
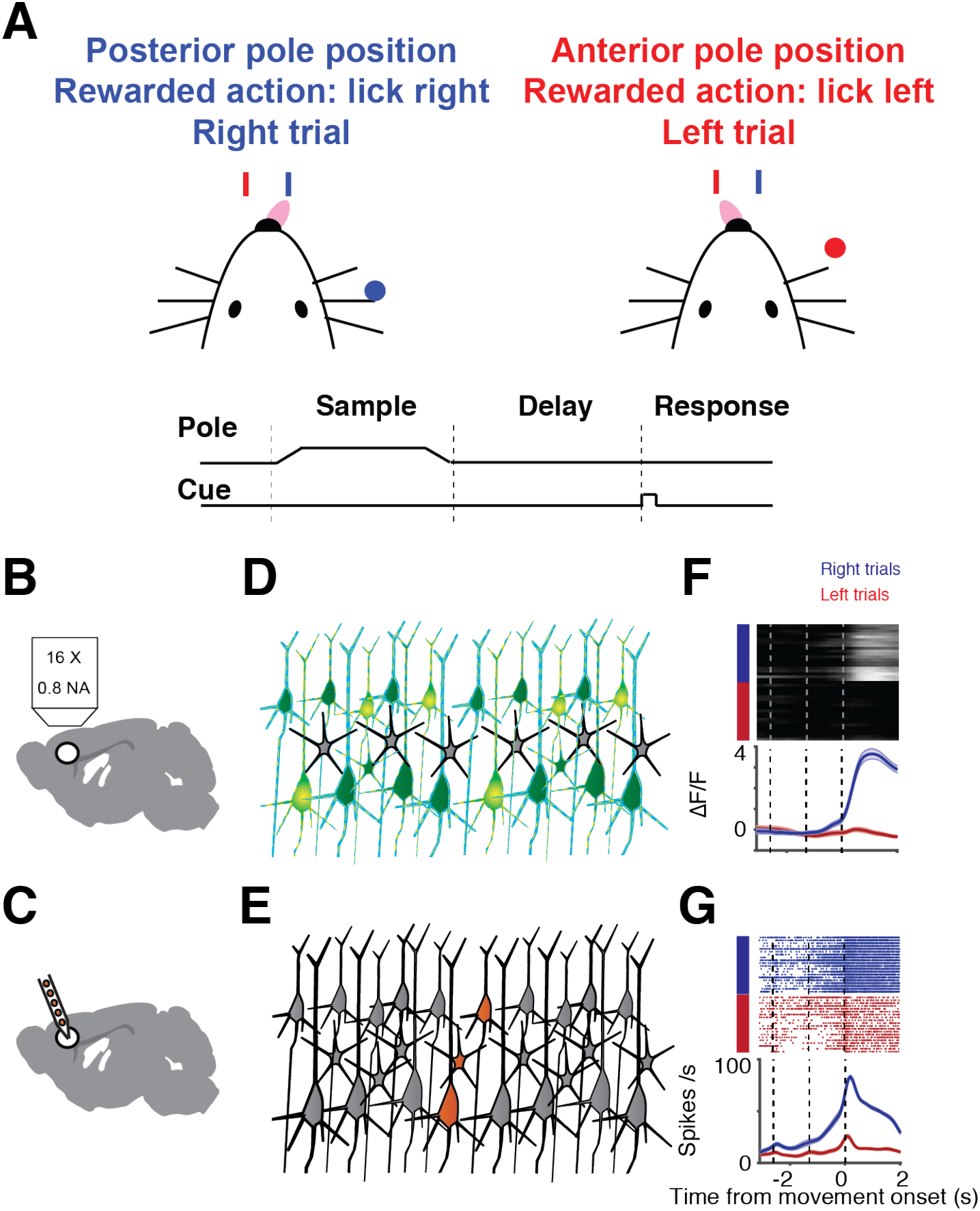
Illustration of sampling population activity in anterior lateral motor cortex using imaging and electrophysiology. **A.** Delayed-response, two alternative forced-choice task. Mice discriminated a pole position (anterior or posterior) and reported it by directional licking (lick right, blue; lick left, red) after a delay period. End of delay period was signaled by an auditory cue **B.** Schematic of imaging setup. **C.** Schematic of electrophysiological setup. **D.** Schematic of neurons sampled by imaging (green). **E.** Schematic of sampled neurons by electrophysiology (orange). **F.** Example neuron, imaging. Top, individual trials (blue, right trial; red, left trial). Bottom, mean activity (mean, thick line; sem., shaded area). **G.** Example neuron, electrophysiology. Top, raster plot. Bottom, peri-stimulus time histogram (PSTH).

Two-photon calcium imaging and ephys were performed in left anterolateral motor cortex (ALM; Figure 1BDF). We report the results of three variants of calcium indicators in this study: GCaMP6s delivered by viral gene transfer, and GCaMP6s and GCaMP6f expressed in Thy-1 transgenic mice. In the first series of imaging experiments, neurons were transduced with adeno-associated virus expressing GCaMP6s (6s-AAV), a widely-used method (Chen et al., 2013; Peron et al., 2015a) (data from (Li et al., 2015), 1493 neurons, 4 mice). In the second, neural activity was recorded by imaging transgenic mice expressing GCaMP6s in cortical pyramidal neurons (6s-TG, data from (Chen et al., 2017), 2293 neurons, 1 mouse). We treated these datasets separately since the mode of delivery of GCaMP can affect its properties. Specifically transgenic GCaMP typically results in neurons that have lower GCaMP6 expression levels and faster fluorescence dynamics compared to neurons transduced with AAV (Dana et al., 2014). Finally, we collected a dataset obtained with a faster, but less sensitive indicator, GCaMP6f (6f-TG, 2672 neurons, 2 mice). We refer to these three datasets as 6s-AAV, 6s-Tg and 6f-Tg, respectively. We compared this data to ephys data acquired with silicon probes that record multiple neurons simultaneously (720 neurons, 19 mice (Li et al., 2015)) (Figure 1CEG). Ephys recordings were subsampled so that their recording depths matched the generally more superficial calcium imaging experiments. Neurons were recorded by 6s-AAV and 6s-Tg at 120 – 740 µm. The matched ephys subset was taken at 100 – 800 µm leaving n = 720 neurons. Neurons were recorded by 6f-Tg at 140 – 470 µm. The matched ephys subset was taken at 100 – 470 µm, leaving n = 225 neurons.

### Filtering of selectivity by calcium imaging

Individual ALM neurons exhibit diverse temporal dynamics, including changes in selectivity over time (Figure 2B) (Guo et al., 2014b; Li et al., 2015; Wei et al., 2019). We classified dynamics into three categories: ‘monophasic’ neurons showed consistent selectivity across the trial (Figure 2A); ‘multiphasic’ neurons changed selectivity over time (defined as having consistent selectivity for at least 335 ms which then changes and remains stable for at least 335 ms more) (Figure 2B); ‘non-selective’ neurons responded similarly across trial types but were still modulated during the task (Figure 2C). The proportion of monophasic selective neurons was similar between the datasets (58% ephys; 66% 6s-AAV; 50% 6s-Tg; 45% 6f-Tg). However, the ephys data set contained a substantial proportion of multiphasic neurons (220/720; 31%), much larger than the imaging datasets (6s-AAV: 76/1493, 5%; 6s-Tg, 98/2293, 4%; compare to matched ephys, 220/720, 31%; 6f-Tg, 69/2672, 3%; compare to matched ephys, 52/225, 20%; p < .001, *χ*^2^ test; Figure 2D-F). As neural response properties can change across cortical layers, we performed a more detailed analysis of the effect of recording depth on single neuron selectivity and find that selectivity was reduced in imaging compared to ephys across depths (Figure S1A-D).

**Figure 2.**
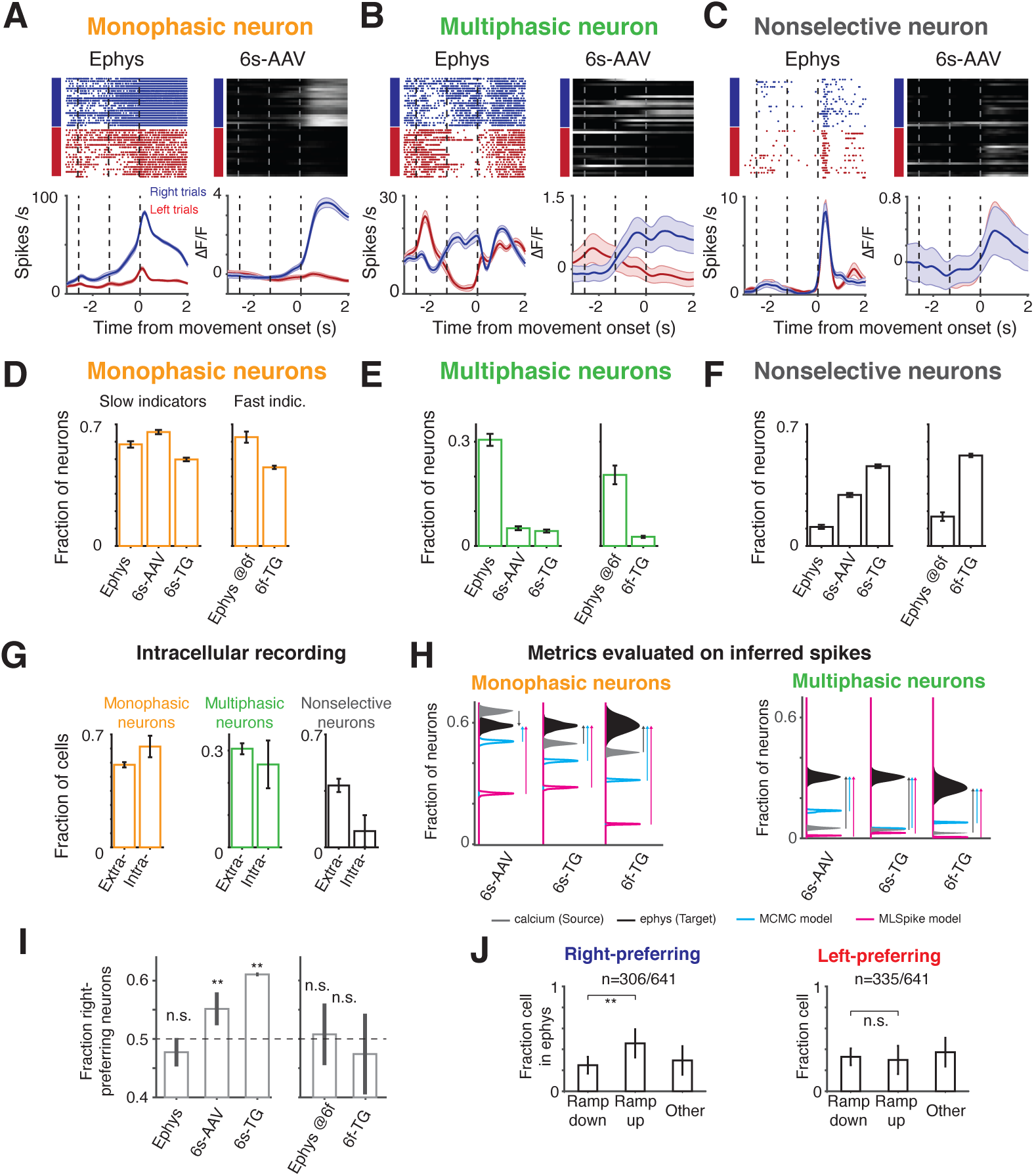
Single neuron trial-type selectivity differs between imaging and ephys. **A.** Example neurons with monophasic selectivity. Left, ephys; right, imaging. **B**, Same as A for multiphasic neurons. **C**, Same as A for non-selective neurons. **D-F**, Fraction of selective neurons in depth-matched ephys (“ephys @6f” indicates depth matched to the more superficial 6f-TG recordings) and when imaged with 6s-AAV, 6s-TG, or 6f-TG. **D.** Fractions of monophasic neurons. **E.** Fraction of multiphasic neurons. **F.** Fraction of nonselective **G**. Proportion of multiphasic neurons in intracellular recordings is similar to that in extracellular recordings. Bar shows fraction of neurons in each of the categories for extracellular (left) and intracellular (right) ephys. **H**. Effect of spike inference on estimates of fractions of monophasic (left) and multiphasic (right) neurons. The distribution of fraction of neurons for imaging data (source data), is given in gray for 6s-AAV (top), 6s-TG (middle) and 6f-TG (bottom). The distribution for ephys (target data) is in black. Distributions from inferred spike rates from MCMC (Pnevmatikakis et al., 2013) are in cyan and for MLSpike (Deneux et al., 2016) are in magenta. Arrows denote the difference between the imaging data and ephys data (gray arrow) or inferred ephys and ephys data (cyan arrow for MCMC and magenta arrow for MLSpike). **I.** Fraction of right-preferring neurons in the different datasets divided into slow indicators (left) and fast indicators (right). **J.** Bar plot of fractions of ramp-down, ramp-up and ‘other’ cells in ephys for right-preferring (left) and left-preferring neurons (right).

What could account for this difference? Ephys records a sparse subset of neurons blindly, which could introduce biases, for example a bias towards neurons with higher spike rates (Figure S1E-I). In contrast, in our imaging experiments all visualized neurons were analyzed. In addition, the spike sorting procedure used to identify units from raw electrode potentials can introduce artifacts, including erroneous merging of neurons (Figure S1J). However, we found that these factors were unlikely to explain our results for two reasons. First, we considered intracellular recordings for which spike sorting is not required (Guo et al., 2017). The fraction of multiphasic neurons was 25.7% (n = 9/35), similar to the extracellular ephys data (p = .67, *χ*^2^ test; Figure 2G) but significantly different from the imaging data (p < .001, *χ*^2^ test with both 6s-AAV and 6s-Tg). Second, we tested the question of spike-sorting induced biases by considering synthetic data in which we deliberately introduced merges at different probabilities. Merging neurons generated more multi-selective neurons when two neurons with different temporal selectivity profiles were merged, but the ratio of accidental merging had to be unrealistically high to explain the difference between datasets (Figure S1J).

Another source of difference could be that our comparisons so far were performed on the imaging data, not on spike rates inferred from the imaging data. Spike inference algorithms attempt to undo the transformation from spikes to claicum-dependent fluorescence, thereby recovering spike times (or spike rates) from imaging data (Berens et al., 2018; Deneux et al., 2016; Pnevmatikakis et al., 2013; Pnevmatikakis et al., 2016; Theis et al., 2016; Vogelstein et al., 2010; Vogelstein et al., 2009). We tested two published methods: MLSpike (Deneux et al., 2016) and MCMC (Pnevmatikakis et al., 2013). In our hands, spike inference only partially corrected the differences between the datasets and in some cases actually pushed the data even further apart (Figure 2H). For instance, MLSpike produced even lower proportions of multiphasic neurons. MCMC was more accurate, increasing the proportions of multiphasic neurons, but still far short of the actual proportion in the ephys dataset (and for 6s-Tg and 6f-Tg decreased instead of increased the proportion of monophasic neurons). Deconvolution at best recovered about half of the missing multiphasic selectivity (6s-AAV, 18%; 6s-TG, 17%, compared to 31% in matched ephys; 6f-TG, 8%, compared to 20% in matched ephys; Figure 2H).

Differences between calcium and ephys were not limited to the temporal nature of responses but were also present in trial-type selectivity. Namely we found that in the ephys dataset right-preferring neurons (i.e., neurons whose firing rate before right licks was higher than before left licks) were as common as left-preferring neurons (Figure 2I, left; p = .118, *χ*^2^ test) (Guo et al., 2014b; Li et al., 2015). The same was true for imaging with a fast calcium indicator (Figure 2I, right; p = .102, *χ*^2^ test), but not for imaging with slow indicators (Figure 2I, left; p < .001, *χ*^2^ test; Figure S4B, spike-inference measure). What could be the cause of these differences? Spike rates in individual ALM neurons often increase or decrease during a trial in ramp-like patterns (Guo et al., 2014b; Inagaki et al., 2018; Li et al., 2015). Right-preferring selectivity was more often associated with neurons ramping up on right trials, whereas left-preferring selectivity included many neurons with firing rates ramping down in the non-preferred (‘right’) trial (Figure 2J). The large difference between the rise and decay times of calcium indicators could lead to differences in how the selectivity of neurons that ramp up or ramp down gets transformed by the indicator. To test to what degree such explanations explain the data we developed a model of the spike-to-fluorescence transformation.

### Simultaneous loose-seal electrophysiology and calcium imaging

Modeling the spike-to-fluorescence transformation requires simultaneously electrophysiology and calcium imaging at the level of individual neurons. Since this data is not available for the transgenic mice used here we performed loose-seal recordings and calcium imaging in individual neurons (Figure 3). The dataset consists of gCAMP6f- and gCAMP6s-expressing L2/3 neurons in transgenic mice (6s-TG, 22 cells; 6f-TG, 18 cells; **Table S1**). This new data more than doubles the number of previously available simultaneously recorded neurons (Theis et al., 2016). In addition we used published data with AAV-based gene transduction (Chen et al., 2013)(http://dx.doi.org/10.6080/K02R3PMN). Bursts of spikes produced fluorescence transients in the imaged neurons (Figure 3A-D). The ability to detect single spikes varied considerably between neurons (Figures 3E-G, S2). The detection of single spikes was lower in transgenic mice than with AAV-based gene transduction, likely reflecting the lower expression level in the transgenic mice.

**Figure 3.**
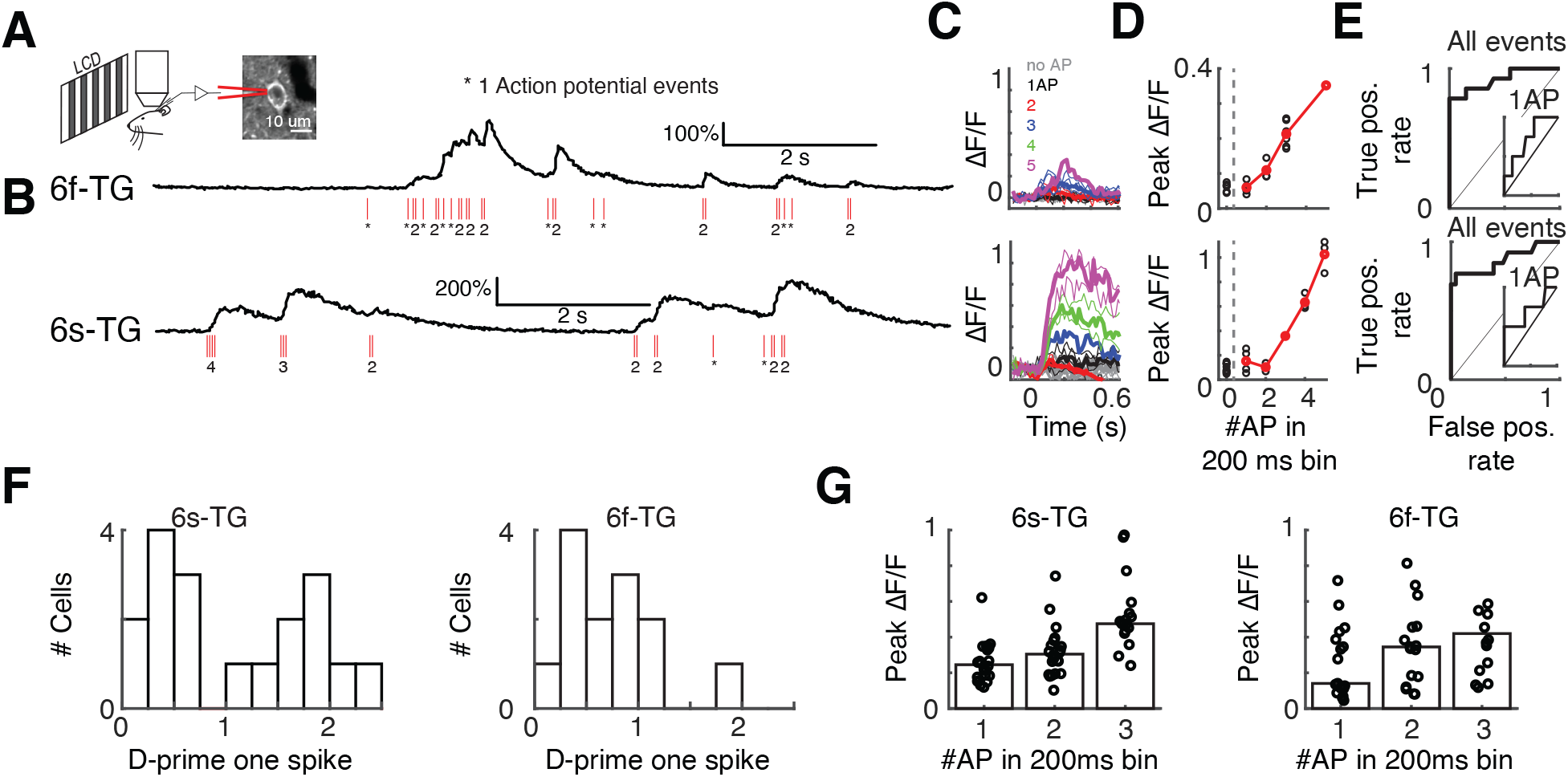
Simultaneous loose-seal recordings and calcium imaging of layer 2/3 pyramidal neurons *in vivo*. **A.** Illustration of the recording setup. Transgenic mice expressing GCaMP6s (GP4.3) or GCaMP6f (GP5.17) were lightly anesthetized and viewed drifting grating visual stimuli. GCaMP-expressing L2/3 neurons were recorded in the loose-seal mode during calcium imaging. **B.** Example recordings from neurons expressing GCaMP6f (top, 6f-TG) and GCaMP6s (bottom, 6s-TG). Red ticks, spikes. **C.** Traces of fluorescence dynamics following different numbers of action potentials (APs) for example neurons. Top, 6f-TG; bottom, 6s-TG. Gray, no AP; black, a single AP; red, 2 APs; blue, 3APs; green, 4APs; magenta, 5APs. Thin lines, single trials; thick lines, average. **D**. Peak fluorescence increases as a function of the number of spikes in 200 ms bins. Black, single trials; red, trial average. **E.** ROC curve of all spike events. Inner panel, ROC curve for single AP events. **F.** Distribution of d-prime for single spikes across cells. Left, 6s-TG; right, 6f-TG. **G.** Mean peak fluorescence changes as a function of number of spikes in 200 ms time intervals across cells. Left, 6s-TG; right, 6f-TG. Each circle corresponds to a recorded neuron. Bars indicate average.

### Spike-to-fluorescence transformations explain differences in single neuron selectivity

Using the newly recorded data we developed a spike-to-fluorescence (S2F) forward model to generate a synthetic calcium imaging data based on a neuron’s spike train (Figure 4A) (Akerboom et al., 2012; Chen et al., 2013; Lütcke et al., 2013; Yasuda et al., 2004). In brief, spike times were first converted to a latent variable, c(t), by convolution with a double-exponential kernel, with parameters rise-time (***τ***_r_) and decay-time (***τ***_d_). This latent variable was pushed through a non-linearity, F(c), with a non-linearity sharpness parameter (k), a half-activation parameter (*c*_1/2_, corresponding to the half-rise point of the nonlinearity) and a maximum fluorescence change (F_m_) (**Materials and methods**). The neurons were well-fit by the model (Figure S3B; variance explained, 6s-AAV, .87 ± .17, mean ± std.; 6s-TG, .80 ± .20; 6f-AAV, .82 ± .27, mean ± std.; 6f-TG, .66 ± .23). The inferred parameter values reflected known indicator kinetics. For instance, the decay times measured for neurons expressing GCaMP6s were longer than those expressing GCaMP6f (Figure 4B). However, there was substantial variability between the parameter values inferred across neurons (Figures 4B, S3). This variability is one factor that could explain the difficulty of the inversion of calcium responses which is central to spike inference approaches. We refer to simulations of calcium-dependent fluorescence based on application of the S2F model to spiking activity as ‘∆F/F_Synth_’.

**Figure 4.**
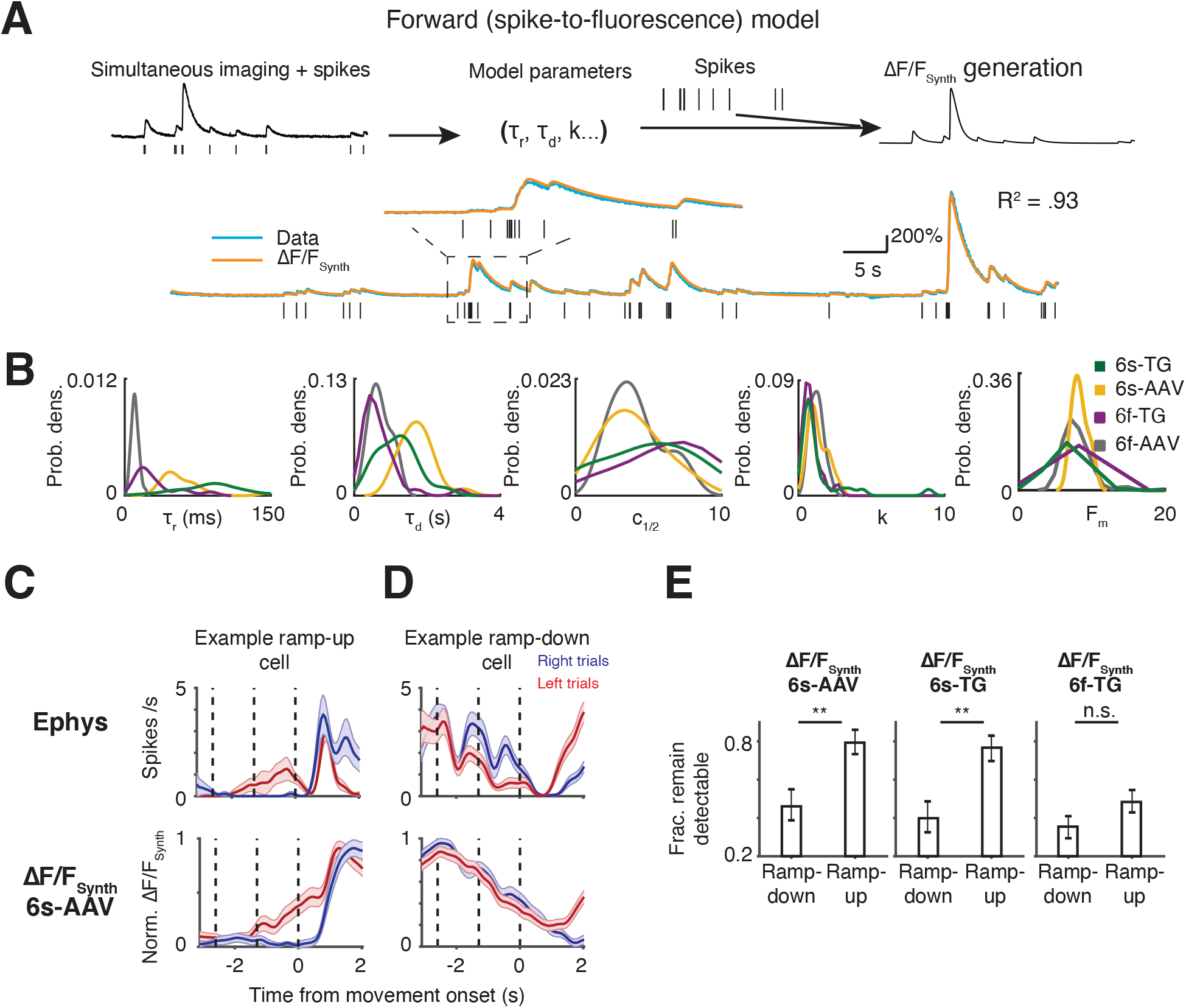
Forward modeling of the spike-to-fluorescence transformation largely explains difference in selectivity patterns. **A.** spike-to-fluorescence model. Top: schematic plot of the spike-to-fluorescence (S2F) forward model that generates a synthetic fluorescence trace (∆F/F_Synth_) from an input spike train. Middle: example fit and data. Experimental, measured ∆F/F (blue) is overlaid with the simulated ∆F/F_Synth_ (orange) from the S2F model. The input to the model, the simultaneously recorded spikes (black), is shown below the traces. **B**. Distributions of the inferred model parameters for different indicators (yellow: 6s-AAV; green: 6s-TG; Purple: 6f-TG; gray: 6f-AAV. **C.** An example ramp-up cell (top, ephys; bottom, 6s-AAV synthetic); selectivity remains detectable in synthetic imaging data. **D.** An example ramp-down cell (top, ephys; bottom, 6s-AAV synthetic); selectivity becomes undetectable in synthetic imaging. **E.** S2F model predicts that selectivity of ramp-down neurons but not ramp-up neurons, would be often obscured in imaging datasets. Bar plot shows fraction of cells that remain detectably selective in synthetic imaging (6s-AAV synthetic, left; 6s-TG synthetic, middle; 6f-TG synthetic, right) plotted separately for ramp-down and ramp-up cells.

We applied the model to ramp-up and ramp-down neurons. For ramp-up cells the separation of activity across trial types was retained in ∆F/F_Synth_, albeit with slower dynamics (Figure 4C). In contrast, for many ramp-down cells ∆F/F_Synth_ became non-selective (Figure 4D). Overall, selectivity was conserved more frequently for ramp-up cells than for ramp-down cells. Since right-preferring cells were more often associated with ramp-up dynamics, and calcium imaging is more likely to capture ramp-up selectivity than ramp-down selectivity, the model explains the greater fraction of right-preferring neurons in the calcium imaging data (Figure 4E). This was true whether a neuron happened to be a right- or left-preferring neuron, i.e., there were no significant differences in the fraction of detectability in the synthetic data once the data was broken down into two categories, ramp-up and ramp-down (p > .05, *χ*^2^ test for all imaging conditions; Figure S4A). Consistent with the difference being produced by the slow decay kinetics of GCaMP6s, there was little difference between the fraction of right- and left-preferring neurons in the 6f-TG data (p > .05 for both cell types). In line with these results, we found that the forward model accounted for the drop in multiphasic neurons presented in the previous section (Figure S4C). These data show that the spike-to-fluorescence transformation introduces systematic discrepancies in comparing the same analysis performed on ephys or imaging data.

### Dimensionality reduction emphasizes different sources of variance in ephys and imaging

Large-scale recording methods are often used in combination with dimensionality reduction techniques to provide a compact description of the data (Cunningham and Yu, 2014). For example, principal component analysis (PCA) finds modes of population activity that capture the largest amount of variance in neural activity (Cunningham and Yu, 2014). Data visualization and analysis are often performed after truncating the decomposition after a few components. We found that the contribution of different sources of variance to the first principal components diverges between ephys and imaging. Accordingly, truncation of PCA in the first few principal components can lead to a qualitatively different PCA decomposition of neural activity between ephys and imaging.

We found substantial differences in performing PCA on ephys and imaging datasets. First, the content of the first PCs was remarkably different between ephys and imaging. In the ephys data, variance in the first PC was mostly due to temporal dynamics (98.71 ± 0.06%, mean ± std., bootstrap analysis). In contrast, for GCaMP6s imaging trial-type selectivity was the dominant source of variance in the first PC (6s-AAV: 60.39 ± 0.29%; 6s-TG: 44.51 ± 0.65%) (Figure 5A). This difference was consistent with the temporal smoothing imposed by slower indicators, and as expected temporal dynamics were predominant in the first PC of GCaMP6f, closer to the values found in ephys, (6f-TG: 64.87 ± 2.47%; depth matched ephys: 91.02 ± 0.14%). Second, in the ephys data, a relatively large number of PCs (> 10) contribute substantially to the variance, whereas in imaging and synthetic imaging most variance was explained by the first few PCs (test for number of PCs required to explain 90% of the variance, p < .001; t test, bootstrap).

**Figure 5.**
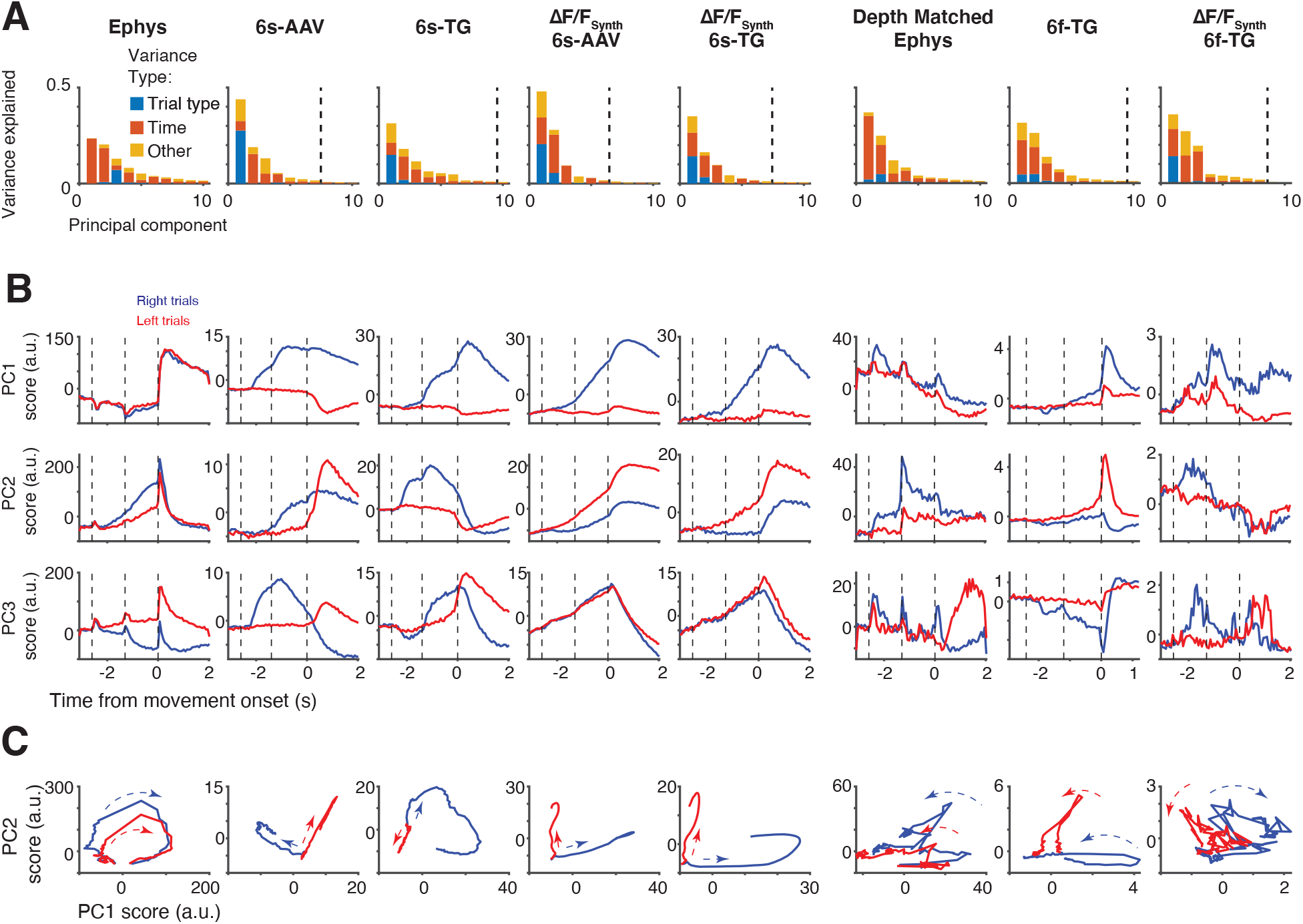
Different sources of variability extracted in dimensionality reduction on imaging and ephys. **A.** Fraction of variance of neural activity explained by principal components 1-10 divided into different sources of variability: red: temporal dynamics; blue: trial type; yellow: other (interaction term). From left to right: ephys, 6s-AAV, 6s-TG, 6s-AAV synthetic, 6s-TG synthetic; ephys depth-matched to 6f-TG recordings, 6f-TG, 6f-TG synthetic. Vertical dashed line indicates the PC index at which the remaining components capture <1% of total variance. **B.** Trial-averaged scores of first three PCs over time (from top to bottom), averaged separately for the two trial types (right trial, blue; left trial, red). Same order from left to right as in A. **C.** Trial dynamics in the first two-PC subspace for the two trial types (right trial, blue; left trial, red). Same order from left to right as in A.

These differences in the sources of explained variance can be seen in the profiles of the PC scores (Figure 5B) as well as in the profiles obtained by a standard exploratory visualization, depicting the evolution of activity over time as a trajectory in the space of the first two PCs (Figure 5C). Spike inference algorithms correctly reduced the amount of trial-type variance in the first principal components (although not fully), but with the caveat that the fraction of variance in the first two principal components was reduced too much (Figure S5).The spike-to-fluorescence model captured the qualitative differences between ephys and imaging, but overestimated the increase in variance in the first two principal components (Figure 5A-C).

### Population activity history affects instantaneous decoding differently in ephys and imaging

Decoding analysis relating population activity to behavioral variables is widely used in systems neuroscience (Ganguly and Carmena, 2009; Harvey et al., 2012; Huber et al., 2012; Wei et al., 2019). Such analyses typically relate the state of population activity at a given time point to a behavioral variable of interest, such as behavioral choice. We performed decoding analysis to predict either trial type or the current behavioral epoch from population activity. Decodability of trial type in ephys increased earlier (one-tail t-test, p < .001), but saturated at a lower level (one-tail t-test, p < .001) than in calcium imaging (Figure 6A). Spike inference models, the MCMC framework in particular, partially reduced the delay of the rise of decodability but overestimated the decrease in decodability yielding lower performance in delay-response epoch than the ephys data (Figure S6). Both observations were recapitulated by the S2F model (delay: one-tail rank sum test, 6s-AAV, p < .001, 6s-TG, p < .001, 6f-TG, p < .001; enhancement: 6s-AAV, p < .001, 6s-TG, p < .001, 6f-TG, p < .001; Figure 6B-C). The counterintuitive result of higher decoding accuracy in imaging for matched population size is explained by the long decay time of slow calcium imaging. The long integration in calcium imaging causes instantaneous decoding on imaging to be equivalent not to instantaneous decoding on spiking data, but to decoding on a more time averaged variable. Such a choice is advantageous when a larger proportion of the selectivity is stable, as was the case in ALM sample and delay selectivity (Li et al., 2016). Consistently, decoders built on ephys that incorporated a one second integration time were more accurate than instantaneous ephys decoders and as accurate as slow indicators (Figure 6D). The delayed increase of decodability was also explained by the forward model. GCaMP6f, with its reduced signal to noise, yielded less accurate population decoders (Figure 6E; spike inference measure, Figure S6).

**Figure 6.**
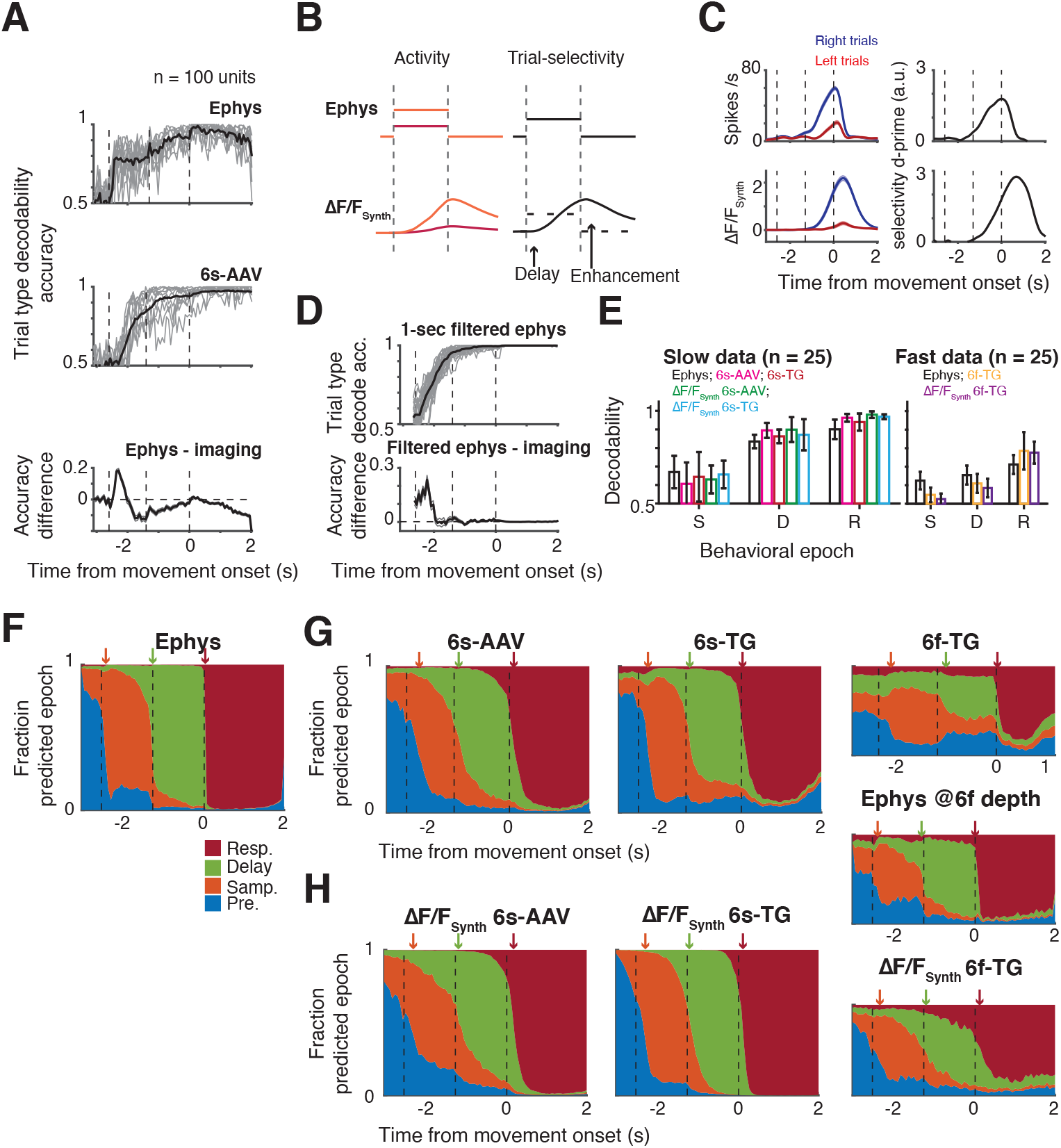
Population decoding differs in sensitivity and temporal profile between imaging and ephys. **A.** Performance of instantaneous regularized linear-discriminant-analysis (LDA) trail-type decoder for 100-unit subpopulations. Vertical dotted lines indicate behavioral epochs, from left to right: presample, sample, delay, response. Top, decoders trained on ephys; middle, decoders trained on 6s-AAV; bottom, difference between the two. For top and bottom plots: individual gray lines show single subsample performance and black thick line shows average. In bottom plot mean is indicated by think line and shaded area corresponds to standard deviation. **B.** Toy model demonstrating observed delayed but enhanced decodability in imaging data. Schematic of relation between activity (left) and decodability (right) when the model has two constant levels of activation for the two trial types (orange and red). **C.** Example cell showing similar behavior to the toy model. **D.** Comparison of decodability from imaging to decodability from 1-second filtered ephys. Top, 1-second filtered ephys; bottom, difference between filtered ephys and imaging. **E.** Comparison of decodability of trial type per behavioral epoch. Decodability for all datasets separated into slow indicators (left) and fast indicators (right). Bars color coded according to dataset. Left: black, ephys; magenta, 6s-AAV; red, 6s-TG; green, 6s-AAV synthetic; cyan, 6s-TG synthetic. Right: black, ephys (depth matched to 6f-TG); orange, 6f-TG; purple, 6f-TG synthetic. **F-G.** Performance of behavioral-epoch LDA decoders. **F.** Probability of decoder based on ephys to assign population activity to each of the different epochs shown in the following color scheme: pre-sample (blue), sample (orange), delay (green), and response (red) epoch; arrows indicate the inferred transition times of epochs from neural codes. **G.** Same plot format as F, but for imaging. **H.** Sample plot format as F, but for synthetic imaging.

For different decoding analyses such averaging can reduce accuracy. For instance, the neurons that can be used to decode trial-type change substantially between the delay and response period, i.e., the patterns of population selectivity are typically dynamical themselves. To test the interaction of these dynamics with calcium indicators, we trained decoders to distinguish the current epoch in the task from the pattern of neural activity. In ephys (Figure 6F) we observed a rapid decrease of the probability of activity to belong to the previous epoch following a change in behavioral epoch, along with a sharp increase in the probability of belonging to the current epoch. In contrast, in the calcium imaging data such changes tended to be delayed and gradual, even for the fast calcium indicator (Figure 6G). This effect was also recapitulated in the synthetic calcium data from the S2F model (Figure 6H). In other words, at the change of a behavioral epoch, the asymmetry of fast rise times and long decay times in calcium indicators yields calcium imaging signals that are a mix of the decaying profile of activity in the previous epoch and the newly activated profile of activity elicited by the response epoch.

### Population dynamics is temporally dispersed in calcium imaging

Neurons show temporally complex responses, even in simple trial-based behaviors (Brody et al., 2003; Li et al., 2015). These spike rate changes are critical for an understanding of neural circuit models of neural computation. Our analysis revealed a qualitative difference in the dynamics between populations recorded by ephys or imaging: a dispersion of the apparent dynamics. That is, the spike rates recorded in ALM peaked at transitions between behavioral epochs (Figure 7A) (Li et al., 2015). In contrast, in the calcium imaging data, peaks of fluorescence were delayed and jittered, causing a more sequence-like appearance (Figure 7B) (Harvey et al., 2012; Scott et al., 2017).

**Figure 7.**
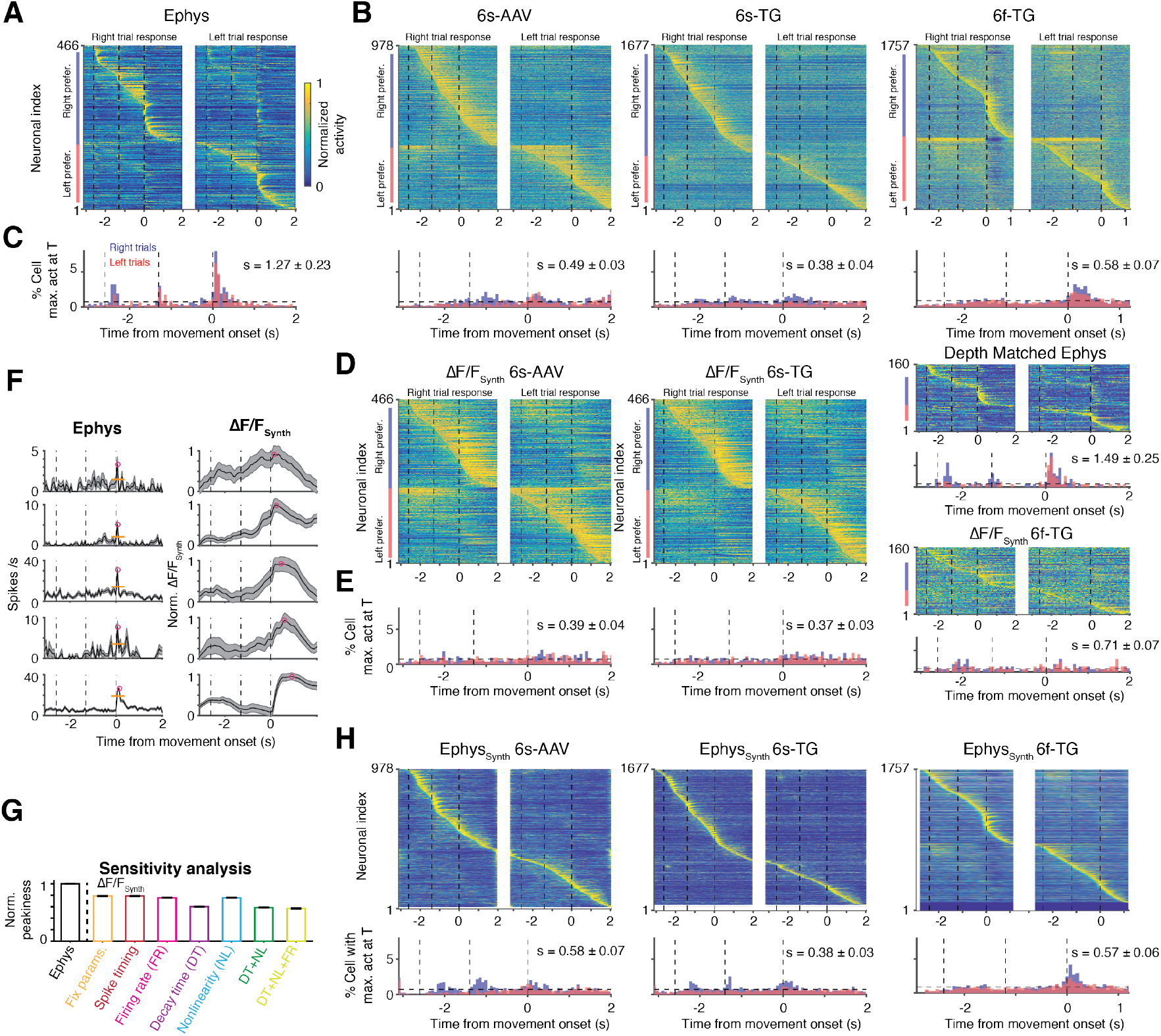
Temporal dispersion of population dynamics differs between imaging and ephys. **A.** Heatmap of normalized trial-averaged firing rates for right trials (left) and left trials (right) for ephys data. Firing rates were normalized to maximum of activity across both conditions. Neurons were first divided into two groups by their preferred trial type then sorted by latency of peak activity. **B.** Same plots as **A** but for 6s-AAV (left), 6s-TG (middle) and 6f-TG (right). Below the 6f-TG are neurons from ephys depth matched to 6f-TG. **C.** Fraction of neurons with a peak at given time point over time. Distribution in time plotted simultaneously for both trial types (red: right trials, blue: left trials, black horizontal line: uniform distribution). Datasets shown left to right (from left: ephys, 6s-AAV, 6s-TG, and 6f-TG respectively). **D-E**. The same plots as B-C for synthetic imaging (6s-AAV synthetic, left; 6s-TG synthetic, middle; 6f-TG synthetic, right). **F.** Example cells with peaks at a similar time in ephys (left; mean activity, thick black line; sem, shaded area; peak, magenta circle; baseline, orange thin line) along with the corresponding synthetic data (right). Neurons are sorted according to their peak times in synthetic imaging (early to late, from top to bottom). **G.** Sensitivity analysis of peakiness by synthetic, artificial data (Materials and methods). Bars show normalized peakiness for the different model variants: (1) identical S2F parameters and identical spike times; (2) identical S2F parameters, jittered spike times (3) identical S2F parameters, variable firing rate (4) identical S2F parameters except for the decay time constant of the calcium indicator that was randomly sampled from its distribution; (5) identical S2F parameters, except for the nonlinearity of the calcium indicator that was randomly sampled from its distribution; (6) both decay time constant and nonlinearity of calcium indicator randomly sampled; (7) variable decay time constant, non-linearity and firing rates. **H**. Same plots as B-C for inferred firing rates from imaging, i.e., synthetic ephys.

To quantify this effect we computed a measure of the ‘peakiness’ of the distribution of neuronal activity (‘s’) across recording modalities as the difference between observed neural activity and temporally uniformly distributed neural activity 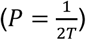:

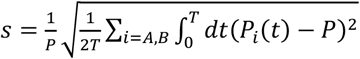

*s* was much larger for the ephys dataset (1.27 ± 0.23) compared to the 6s-AAV (0.49 ± 0.03; one-tail rank sum test, p < .001), 6s-TG (0.38 ± 0.04; one-tail rank sum test, p < .001), and 6f-TG (0.58 ± 0.07; one-tail rank sum test, p < .001) imaging data (Figure 7C). The forward model was able to recapitulate the differences between ephys and imaging (s = 0.39 ± 0.04, 6s-AAV ∆F/F_Synth_; s = 0.37 ± 0.03, 6s-TG ∆F/F_Synth_; s = 0.71 ± 0.07, 6f-TG ∆F/F_Synth_; Figure 7DE). Using the forward model we found that the degree of delay in the peak response is dependent on interactions between multiple factors including the assumed temporal and non-linear parameters of the indicator, as well as the absolute value of the underlying firing rate (Figure 7FG). Here, spike inference algorithms were able to partially undo the difference between imaging and ephys, yielding a reduction in the temporal dispersal (Figure 7H). Similar overall results were obtained with different metrics for the sharpness of the maximum-activity-time distribution relative to a uniform distribution, such as the Kullback-Leibler divergence.

Similar analyses on single neuron and population activity properties were performed on ephys and imaging data from the primary somatosensory cortex with qualitatively similar results (Figure S7).

## Discussion

Calcium imaging using fluorescent protein sensors is a powerful method for recording activity in large neuronal populations (Peron et al., 2015a). In systems neuroscience, cellular calcium imaging fills a complementary role to extracellular electrophysiology. Imaging can sample neural activity densely (Peron et al., 2015a; Peron et al., 2015b) and reveal spatial relationships between neurons with related activity patterns (Kerlin et al., 2010; Ohki et al., 2005). Imaging can be used in a cell-type specific mode to sample rare neuronal populations that are difficult to target using electrophysiology (Fu et al., 2014). Imaging can be combined with post-experiment molecular analysis (Kerlin et al., 2010; Lovett-Barron et al., 2017; O’Connor et al., 2010) or even serial electron microscopy reconstruction (Bock et al., 2011; Briggman et al., 2011). Imaging can track the activity of individual neurons over long time scales to explore the circuit basis of learning (Huber et al., 2012; Komiyama et al., 2010). Finally, imaging allows recording activity in neuronal microcompartments that are not accessible to electrophysiology (Chen et al., 2013; Jia et al., 2010; Petreanu et al., 2012; Xu et al., 2012). Electrophysiological recordings report neural activity with high temporal precision but have limitations of their own. Ephys recordings have a bias towards large neurons with high spike-rates. In addition, the process of transforming raw recordings into spike times associated with individual isolated units, i.e., spike sorting, can introduce artifacts such as merging spikes from different neurons.

Although calcium imaging and ephys are often used almost interchangeably, the quantitative effects of the differences between ephys and imaging on measures typically used in system neuroscience have not been evaluated in a systematic manner. By comparing activity recorded with electrophysiology or imaging from the same circuit during the same behavioral task we showed that the different recording methods can lead to diverging results. On the level of single neurons, the proportion of neurons with specific response properties and different dynamics of selectivity differs between calcium imaging and ehpys. At the level of neuronal populations, we find diverging results for the content of population activity variance (trial condition differences being the main source of variance in imaging while temporal dynamics are the main source of variance in ephys), the relation of population activity to behavior, and the overall pattern of population dynamics. Spike inference algorithms only partially recovered the difference between ephys and imaging across the multiple metrics considered in this study (Figure S8B). Notably, we find large neuron-to-neuron variability in the inferred parameters of a forward, spike-to-fluorescence model. Such variability coupled with the large heterogeneity in firing rates and temporal patterns makes correctly solving the inverse problem difficult, which potentially explains our results. At the same time, most of the differences we found between ephys and imaging were explainable by a forward-model that generates a synthetic imaging experiment counterpart of a neuron’s ephys responses. Such a model takes into account the specific heterogeneity found in ephys recordings and can take into account neuron-to-neuron variability in calcium imaging properties by sampling randomly from the varying parameters of the spike-to-fluorescence transformation.

### Im-phys.org – a website for more detailed comparison ephys and imaging

We presented an extensive dataset with three calcium indicators, extracellular and intracellular electrophysiology and multiple models. However, a research paper still represents a small distillation of all possible analyses. We developed an online resource, im-phys.org (http://im-phys.org/ Figure S8) with three goals. First, the website allows analysis of all combinations of dataset and model, to evaluate the scenario that is most relevant to particular experiments. Im-phys.org allows spike inference algorithms to be systematically tested in real use case scenarios, i.e., not just testing recovery of any aspect of the patterns of spike rates but rather testing the impact of performing spike inference on undoing differences in specific metrics extracted from ephys and imaging (Figure S8B). Second, we hope that other groups will share data, models and analyses to allow more general comparison of ephys and imaging data. Im-phys.org allows submission of data that can be incorporated into various comparisons that are displayed on the website, controlled through UIs. Though few labs have matched ephys and imaging datasets, many labs have one or the either. Our resource can serve to aggregate and combine these datasets, as well as find a best match from an imaging to ephys dataset (Figure S8A). Third, im-phys.org is linked to a github repository containing the analyses code, models (S2F and F2S), and related data. These allow to use or analyses and models on data without sharing it through im-phys.org

### Differences between interrogating population activity by ephys and imaging affect data-driven models

Differences in metrics of population activity between calcium imaging and ephys not only complicate the research literature but can result in the divergence of models used to understand the underlying data. Most population models, whether models in which the single units are modeled in more biophysical detail or more abstractly, are still highly reduced in the way they treat population heterogeneity. As such they often rely on dimensionality reduction of the recorded data to define the aspects of population activity the model is meant to capture. We found substantial differences between ephys and imaging data in application of PCA, and the truncation of the data after a few important data components can further amplify differences. In extreme cases one may be left with subsets that differ dramatically across imaging and electrophysiology. The amplification of difference by dimensionality reduction is relevant not just for modeling of the data, but more generally when generic forms of dimensionality reduction, such as PCA, are used early in the analysis pipeline to improve signal-to-noise ratio (which is important given the limited duration of typical behavioral experiments) for subsequent analysis, such as population decoding. Dimensionality reduction can be hard to avoid when analyzing large datasets (Cunningham and Yu, 2014), but can be modified to be less sensitive to known issues.

### Going forward

Going forward, the discrepancies between ephys and calcium imaging can be reduced by improvements in calcium indicators and adjustments to experimental design. Calcium indicators could be improved on multiple fronts. They could be made faster and less nonlinear (Dana et al., 2018; Inoue et al., 2019). In addition, more uniform expression across cells can allow for more aggressive modeling of the nonlinearities that cannot be reduced, especially when coupled with priors on activity profiles derived from large scale electrophysiology. Faster indicators will result in the effect of previous activity history washing away faster, thus reducing effects that are history dependent. Imaging with multiple types of indicators in different experiments might produce additional constraints and help reduce biases. Voltage imaging holds great potential for fast accurate measurement of spiking activity, at least in sparsely labeled neuronal populations (Abdelfattah et al., 2019; Adam et al., 2019). At the level of experimental design, when population activity in a given behavioral epoch involves fixed dynamics, such as settling to a steady state or consistent ramping, longer trial epochs will allow the effect of the previous dynamical state to decay away. Indeed, we found a smaller discrepancy between the number of multiphasic neurons in ephys and 6s-TG data when the behavioral paradigm was adapted to use longer delay epochs. If priors on the profile of activity are known from electrophysiological recordings, an effort that should become easier as neurophysiology probes become more powerful (Jun et al., 2017), data sharing more common, and preprocessing more standardized, fluorescence-to-spike models can assist in evaluating experimental design and its effect on dynamics.

Overall our results highlight the importance of a deeper understanding of the transformation imposed by calcium imaging. The fact that our model was able to reproduce differences between the recording methods suggests that additional data and associated analysis methodology developments could potentially better address quantitative comparisons between analyses of population activity performed from imaging or ephys data. The online resource we built allows researchers to better understand how the discrepancies we observed would be relevant for the circuit and recording method of interest. More quantitative interpretation of calcium imaging and full utilization of all its advantages will require investment in ground-truth data sets and new statistical approaches. We hope this study and our online resource will catalyze this crucial effort.

## Authorship

Z.W, K.S and S.D conceived the project. B.J.L performed simultaneous ephys-imaging recordings. T.W.C and K.D. performed 6s-TG and 6f-TG imaging experiments. Z.W, K.S. and S.D performed model simulations and analyzed the data. Z.W, K.S and S.D wrote the paper with comments from the other authors.

## Acknowledgement

We thank Arseny Finkelstein, Christopher Harvey, Daniel Huber, Aaron Kerlin, Daniel O’Connor and Louis K. Scheffer for comments on the manuscript and Nuo Li for many useful discussions. This work was funded by the Howard Hughes Medical Institute. Simultaneous recordings and imaging experiments were performed with support from the Genetically Encoded Neural Indicator and Effector (GENIE) project. T.W.C is supported by a career development grant (NHRI-ex-105-10509NC) from the Taiwan National Health Research Institute. K.D, K.S and S.D are funded by the Simons foundation collaboration on the global brain SCGB 542969SPI; Z.W is funded by SCGB 542943SPI; S.D is funded by NIH NS104781.

## Competing interests

All authors have no competing interests declared.

## Materials and methods

### Electrophysiological and imaging population activity recordings

Electrophysiological (‘ephys’) or calcium imaging recordings were performed in separate experiments (Li et al., 2015). Mice were trained to perform a delayed version of a tactile discrimination task. Mice reported the position of a pole (anterior or posterior) by directional licking (lick-left or lick-right) after a delay period. The duration of sample and delay epoch was 2.6 s. In ephys, the delay epoch was 1.3 s; in imaging, it was 1.4 s. Trials with early licking were excluded from analysis. Neuronal depths were 100 to 800 um (ephys), 150 – 740 um (6s-AAV), 120 – 640 um (Thy1-GP4.3 mice, 6s-TG), and 140 – 470 um (Thy1-GP5.17 mice, 6f-TG). Only sessions with more than 20 trials for each type (right-trial and left-trial) were included. For imaging data, we performed a post-hoc detection of outliers and removed trials where more than 30% of the time points contain a signal with 3 standard deviations away from median (these outliers relate to baseline fluctuations across trials, and removing them was necessary for variance-based analysis). Neurons were limited to putative pyramidal neurons. These reduced the total number of neurons with sufficient number of trials, yielding 1493, 2293, and 2672 units for 6s-AAV, 6s-TG and 6f-TG imaging, respectively.

We used two sets of data from loose-seal electrophysiological recordings and imaging from GCaMP6-expressing neurons in primary visual cortex. In one set neurons were transduced with 6s-AAV and 6f-AAV (data from (Chen et al., 2013)). In the other set we used 6s-TG and 6f-TG mice (Dana et al., 2014; Lin et al., 2016). More details of all datasets are described at http://im-phys.org/data.

Simultaneous loose-seal recordings and imaging (Figure 3 and Figure S2) was performed as described previously(Chen et al., 2013) (more details at http://im-phys.org/data). GP4.3 and GP5.17 mice (Dana et al., 2014) were lightly anesthetized (0.5% isoflurane). Drifting grating visual stimuli were used to drive activity in the visual cortex. Loose-seal recordings were made through a craniotomy windows over the primary visual cortex. Two-photon imaging and loose-seal, cell-attached recordings were performed simultaneously. We acquired images in both low (284 × 284 um^2^) and high (38 × 38 um^2^) zoom configurations. Extraction of fluorescence transients was as described (Chen et al., 2013).

To analyze the spike-triggered fluorescence changes, we created 1.2-s snippets around action potentials (APs), where a few APs only happened from 200 ms to 400 ms from the onset of each snippet. We computed baseline fluorescence using the snippets without AP in the entire time series. For snippet with APs, we required the fluorescence changes within the first 200 ms (before APs) was around baseline level (Figure 3C, S2A). We computed ROC curve as the probability if the peak fluorescence changes after 1 (insert panels) or many APs (Figure 3E, S2C) can be detected at certain threshold comparing to baseline fluorescence fluctuations. D-prime is computed as 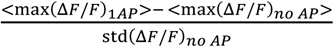.

### Spike-to-fluorescence model

We developed a phenomenological model that converts spike times to synthetic fluorescence time series (Akerboom et al., 2012; Chen et al., 2013; Li et al., 2015; Yasuda et al., 2004). This ‘spike-to-fluorescence’ (S2F) model consists of two steps. First, spikes at times {*t*_*k*_} are converted to a latent variable, c(t), by convolution with a double-exponential kernel:

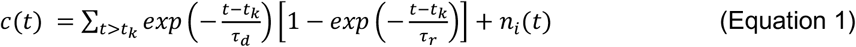

***τ***_r_ and ***τ***_d_ are the rise and decay times, respectively. 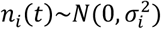 is Gaussian distributed ‘internal’ noise. c(t) was truncated at zero if noise drove it to negative values. Second, c(t) was converted to a synthetic fluorescence signal through a sigmoidal function:

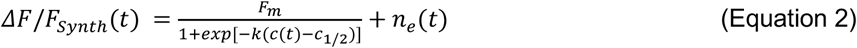

k is a non-linearity sharpness parameter, c_1/2_ is a half-activation parameter, F_m_ is the maximum possible fluorescence change. 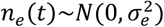 is Gaussian external noise (Maravall et al., 2000; Tsien, 1989; Yasuda et al., 2004).

We estimated the model parameters for each imaging condition using the simultaneous ephys and imaging experiments (Figure S3ABC). We then applied the S2F model to ephys data using parameters randomly sampled from the parameter distributions except for the parameters directly related to the nonlinearity. Since ALM spike rates in ephys vary over a larger range than the spike rates in the primary visual cortex these parameters may be underconstrained. Accordingly, we followed an alternative strategy to choose these parameters for a given neuron. For each neuron, after assigning the rest of the parameters, we transformed the spike trains to calculate the phenomenological calcium variable *c*(t). We then estimated the nonlinear parameters for that neuron by calculating the values that would best transform *c*(t) to the fluorescence dynamics of any neuron in the imaging dataset. For all neurons we were able to find matches with Spearman correlation higher than 0.7 between mean dF/F and mean synthetic dF/F. The parameters inferred in this process recapitulated the correlation structure of c_1/2_ and k found in the data (Figure S3D).

Given the short timeframe over which baseline activity was recorded before each trial started, we extended the pre-trial period by simulating a Poisson spike train for the unrecorded time between trials with a constant rate equal to the baseline mean activity.

To relate this model to previously studied models, Equation 2 can be generalized as *ΔF*/*F*_*Synth*_(*t*) = *f*(*c*(*t*)) + *n*_*e*_(*t*), where *f*(·) and 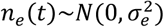 is Gaussian external noise (Maravall et al., 2000; Tsien, 1989; Yasuda et al., 2004). We considered two alternative S2F models used by previous studies (note though that both of these models did not contain internal noise in Equation 1):

#### S2F Linear model

*f*(*c*(*t*)) = *F*_*max*_*c*(*t*) + *F*_0_, where *F*_*max*_ is a scaling parameter (we kept the naming as max to clarify the relationship to other models); *F*_0_ is the baseline (Figure S3G, left).

#### S2F Hill model

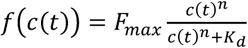, where *F*_*max*_ is the maximum possible fluorescence change; *n* is the nonlinearity; *K*_*d*_ is a half-activation parameter. Model performance is summarized in the supplementary material and reported in http://im-phys.org/analyses for each single cell (Figure S3G, right).

Model parameter sensitivity (Figure S3C) was defined as the decrease of the fraction of explained variance, as a function of the deviation of the parameter value from the estimated solution: 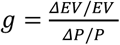, where *P* ∈ {*τ*_*r*_, *τ*_*d*_, *k*, *c*_1/2_, *F*_*m*_}.

### Calcium imaging to spikes for non-simultaneous ephys-imaging recordings

We performed fluorescence-to-spike (F2S) inference using two published models (Deneux et al., 2016; Pnevmatikakis et al., 2013). These were two state-of-the-art generative models based on inference techniques developed by (Deneux et al., 2016; Pnevmatikakis et al., 2013). We used the authors’ code available on GitHub.

### Single neuron analyses

Neural selectivity for left- or right-trials was determined using two-sample t-tests, with neural activity binned over 67 ms, which corresponds to one imaging frame. A neuron was selective if it showed selectivity (p < .05) for >335 ms (5 continuous frames). A selective neuron was multiphasic if the polarity of selectivity switched, with continuous periods of selectivity lasting at least 335 ms long. Selective neurons that were not classified as multiphasic according to this criterion were classified as monophasic.

Selective neurons (mono- and multiphasic) were classified into left- and right-preferring cells according to the condition in which their activity was higher (Figure 2IJ). Ramp-down (ramp-up) were defined as neurons that have activity that is greater (less) in the baseline epoch compared to the delay epoch (paired t-test, p < .05 across trials). Note that ramp-down cells were excluded from the analysis of peakiness (Figure 7).

### Principal component analysis

Principal Component Analysis (PCA) was performed on the activity of neurons averaged across trial type (*s* ∈ {*left, right*}):

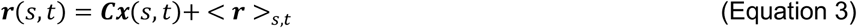

***r*** is a *n* × 2*T* matrix, where *n* is the number of recorded units in each dataset and T is the number of time points for each trial type. < ***r*** >_*s,t*_ is a vector of the mean activity of each neuron across time and trial type. ***x***(*s*, *t*) is an *n* × 2*T* PC score matrix, where the 𝑖th row corresponds to the *i*th PC score. We estimated the relative contribution to each PC of the different forms of variance: temporal dynamics, trial-type selectivity and other. Explained variance (EV) of temporal dynamics *EV*_*i*_(*t*) and trial-type selectivity *EV*_*i*_(*s*) for the *i*th principal component (PC) were computed as:

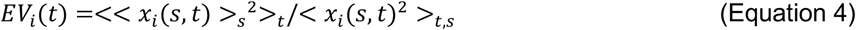

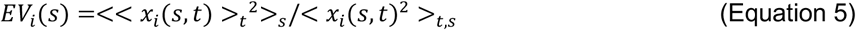

respectively. 6f-Tg related population analyses were only applied to cells with ROC > 0.7.

### Population decoding

We applied regularized linear discriminant analysis (LDA) on neural dynamics grouped into bins corresponding to single imaging frames (67 ms) to compute the instantaneous decodability of trial type. Regularization was performed by sparsity-regularized LDA (Guo et al., 2007; Wei et al., 2019). The optimal LDA decoder was computed separately for each time bin using correct trials only. We estimated performance for the instantaneous LDA decoder by sampling subsets of units and averaging 100 subsamples. We separated the trials of each neuron into non-overlapping training (70%) and testing (30%) sets. The instantaneous decoder of trial type was computed from training set and its performance was evaluated on the testing set.

We tested the ability of neuronal population activity at different times to discriminate the behavioral epoch by using a four-class LDA (Figure 6F-H). We defined the latency of neuronal response to behavioral epoch by the first time at which decoding reached a 0.7 accuracy threshold (arrows on Figure 6F-H). Regularization was performed by sparsity-regularized LDA (Guo et al., 2007; Wei et al., 2019).

### Sensitivity analysis of peakiness

We used as a reference value an artificial, synthetic ephys dataset with 50 neurons whose firing rates were manually set to be non-zero only at the time corresponding to one imaging frame. From left to right in Figure 7G, S2F model was configured (1) using the same parameters for all cells, except that the internal noise and external noise were randomly generated (at the same amplitudes); (2) using the same parameters for all cells, except that the spike times were jittered within the time length of the frame (i.e., all spikes were kept in the same image frame); (3) using the same parameters for all cells, except that the spike rates in the original frame varied from 0.1 Hz to 5 Hz (spike trains generated using Poisson process); (4) using the same parameters for all cells, except that the decay time constant of calcium indicator was randomly sampled from its distribution; (5) using the same parameters for all cells, except that nonlinearity of calcium indicator was randomly sampled from its distribution; (6) both decay time constant and nonlinearity of calcium indicator were randomly sampled; (7) the same as (6) except that the spike rates in the original frame vary from 0.1 Hz to 5 Hz (spike trains generated using Poisson process).

### Distributions of measures

For S2F model, one can randomly sample all the parameters from the distributions measured using simultaneous ephys-imaging recordings and all possible noise levels. The distribution of a measure *ψ* (e.g. fraction of mono-selective neurons, peakiness etc.) can then be computed through synthetic data using randomly sampled S2F models. Specifically:

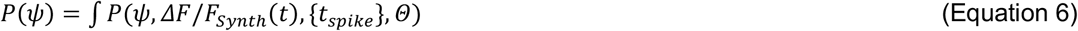

where the joint distribution can be formulated through a chain rule:

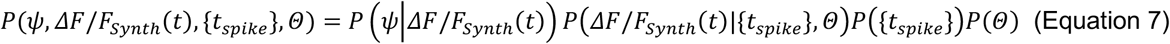

where *P*(*ΔF*/*F*_*Synth*_(*t*)|{*t*_*spike*_}, *Θ*) is derived from Equations 1, 2, and *P*(*ψ*|*ΔF*/*F*_*Synth*_(*t*)) describes probability of measure *ψ* at a given value for dynamics *ΔF*/*F*_*Synth*_(*t*), *P*({*t*_*spike*_}) is the empirical distribution of spike events in ground truth ephys and *P*(*Θ*) is the distributions of S2F parameters.

For unsupervised-learning-based F2S models (i.e. MCMC and MLSpike), we performed 100 subsamples of deconvolved synthetic ephys data to estimate distribution of the parameters.

## Supplementary materials

**Figure S1.**
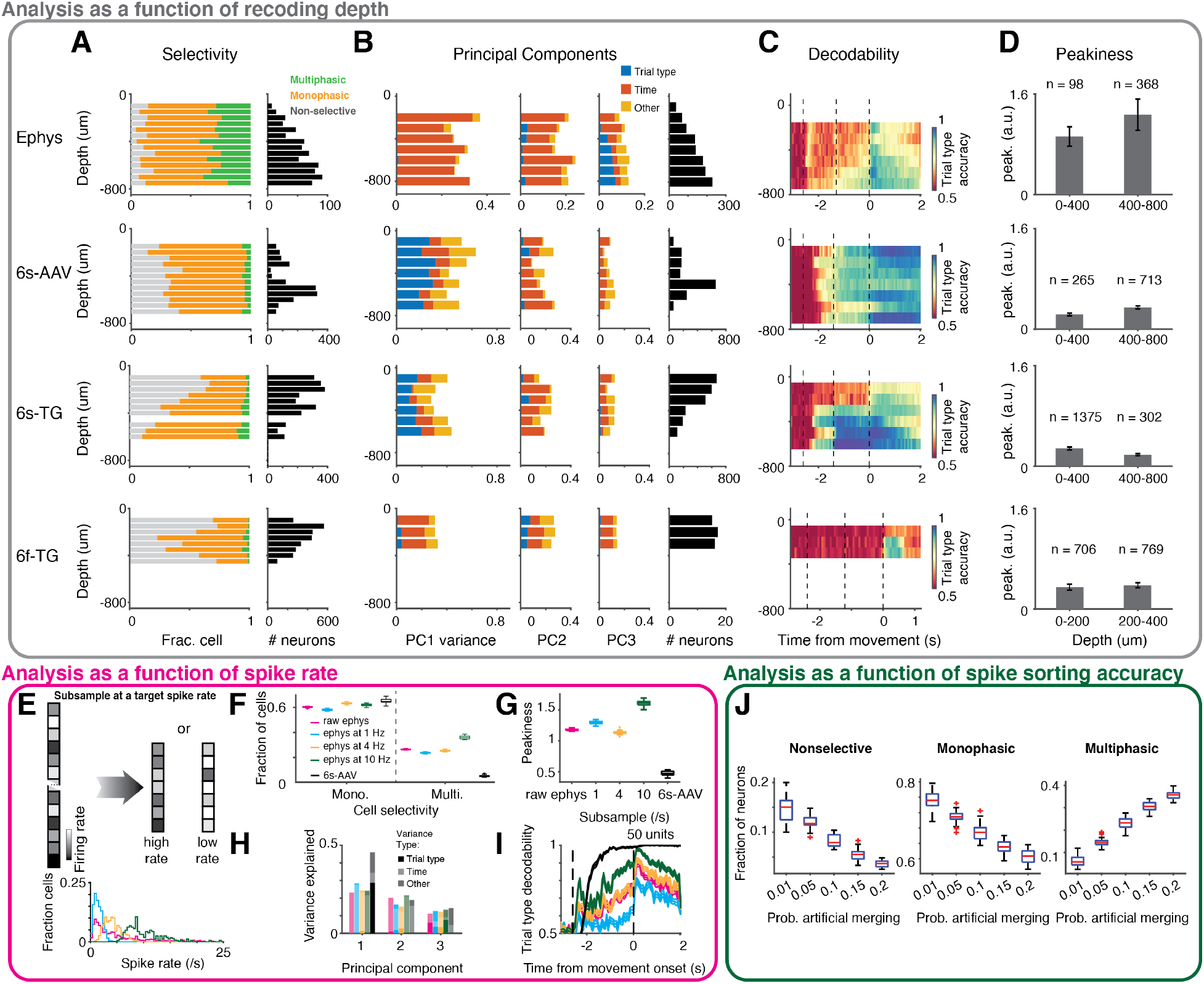
Effect of recording depth and firing rate in ephys. **A-D.** Analysis as a function of recoding depth. **A.** Single neuron selectivity-type analyses. Left: horizontal bar plots show breakdown of population into selectivity types (gray: non-selective neurons, orange: monophasic-selective neurons, green, multiphasic-selective neuron. Right: horizontal bar plot shows number of neurons at each depth. The ratio of monophasic- to multiphasic selective neuron was roughly identical across depths (*χ*^2^-test to depths with n > 50 cells, ephys: p = .19; 6s-AAV: p = .73; 6s-TG: p = .97; 6f-TG: p = .43). For the same depth, ephys has more selective neuron and more multiphasic selective neuron than imaging (*χ*^2^-test, p < .001 for all). **B**. Percentage of variance of neural activity explained by each principal component (Figure 5). Left: length of horizontal bar shows fraction of variance in each principal component. Colors show breakdown into different types of variance (blue: trial-type, red: time, orange: other). Right: horizontal bar shows number of neurons in each depth. For the same depth, the 1^st^ PC show more temporal dynamics content in ephys and 6f-TG (*χ*^2^-test, p < .001 for all), while that show more trial-type content in 6s-AAV and 6s-TG (*χ*^2^-test, p < .001 for all). **C**. Decodability of trial type (Figure 6). The number of cells at each depth is identical to that in PCA analyses. The decodability differs across depths, where the neurons in superficial layers show weak decodability of trial type in sample-delay epoch (multivariate ANOVA test on time-series to depths with n > 50 cells in ephys, 6s-AAV and 6s-TG; that to depth with n > 10 cells at ROC > 0.7 in 6f-TG; p < .001, 1000 bootstrap). For the same depth, the average decodability of trial type is higher in late delay to early response in imaging than that in ephys (rank sum test, p < .001 for all, 1000 bootstrap). **D**. Peakiness (Figure 7). The peakiness differs across depths (rank sum test, p < .001, 1000 bootstrap). For the same depth, peakiness is higher in ephys than imaging (rank sum test, p < .001 for all, 1000 bootstrap). **E-I.** Analysis as a function of spike rates. **E.** Schematic of resampling procedure to target firing rate and distribution of firing rates after subsampling to different average spike rates (magenta: original data; cyan: ephys subsampled to 1 Hz average; yellow: ephys subsampled to 4 Hz average; green: ephys subsampled to 10 Hz average. **F.** Effect of target firing rate subsampling on fraction of monophasic (left) and multiphasic neurons (right) **G.** Values of peakiness are shown with the same color code as F. **H.** Fraction of variance in the first principal components are shown by length of bar with same color code as F. Saturation of bar shows the breakdown into different components of variance (trial-type, time, other). **I.** Trial-type decodability over time shown with the same color code as F and with 6s-AAV added as a reference. **J.** Analysis as a function of spike sorting accuracy -- possible effects of merging. Increased fraction of multiphasic neurons is unlikely to have stemmed exclusively from failures of spike-sorting. Box plots indicate fraction of neurons in each selectivity class (left: non-selective, middle: monophasic, right: multiphasic) as a function of increased probability of artificially induced merging between two neurons. Dashed line indicates fraction of selectivity type found in the ephys dataset.

**Figure S2.**
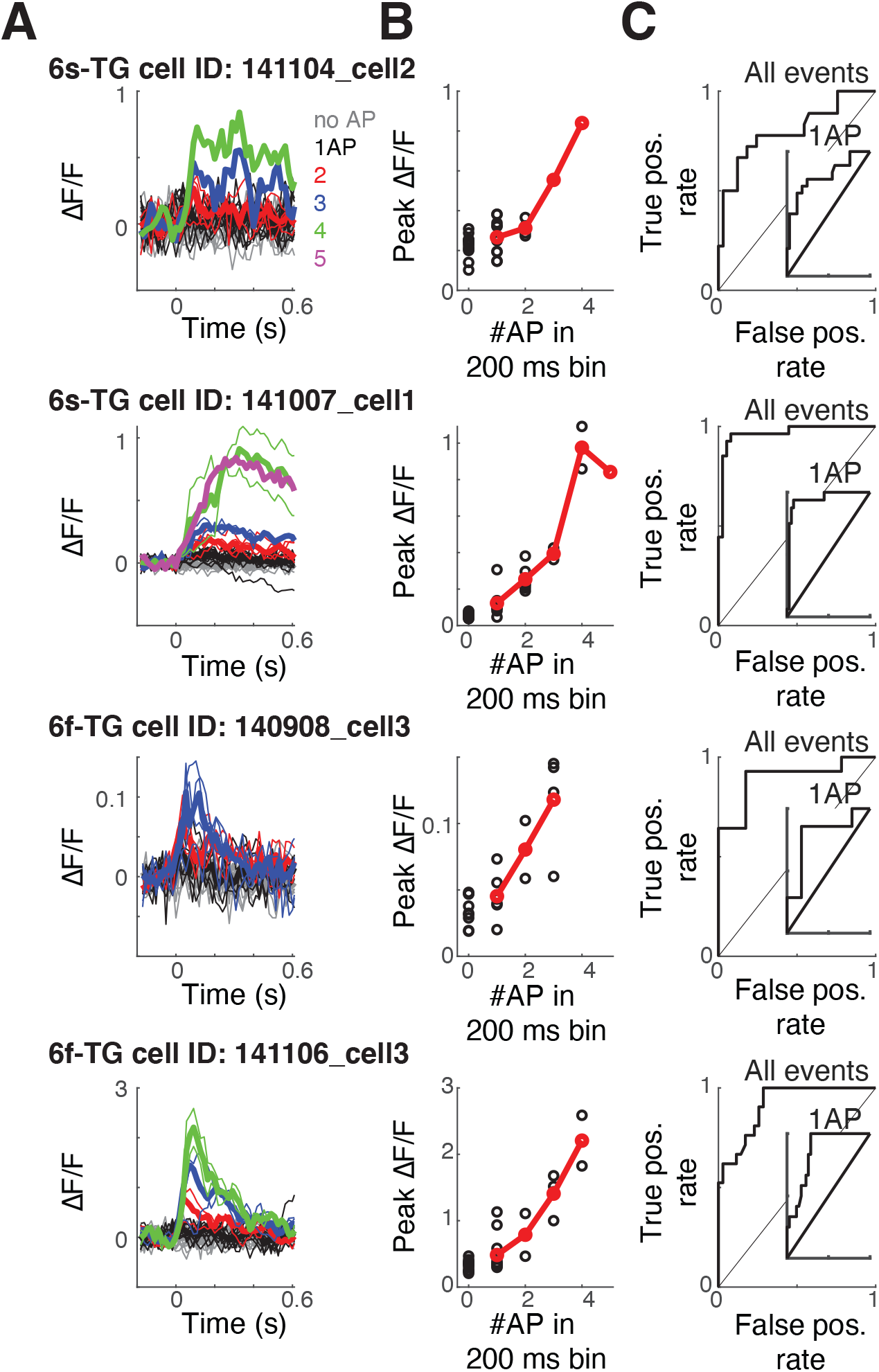
Single- and few-AP responses of neurons in transgenic GCaMP6s and 6f mice. **A.** Traces of fluorescence dynamics following different numbers of action potentials (APs) for example neurons (same plots as Figure 3C for additional examples). Gray, no AP; black, a single AP; red, 2 APs; blue, 3APs; green, 4APs; magenta, 5APs. Thin lines, single trials; thick lines, average. **B**. Peak as a function of the number of spikes (same plots as Figure 3D for additional examples). Black, single trials; red, trial average. **C**. ROC curve of all spike events. Inner panel, ROC curve for single AP events (same plots as Figure 3E for additional examples).

**Figure S3.**
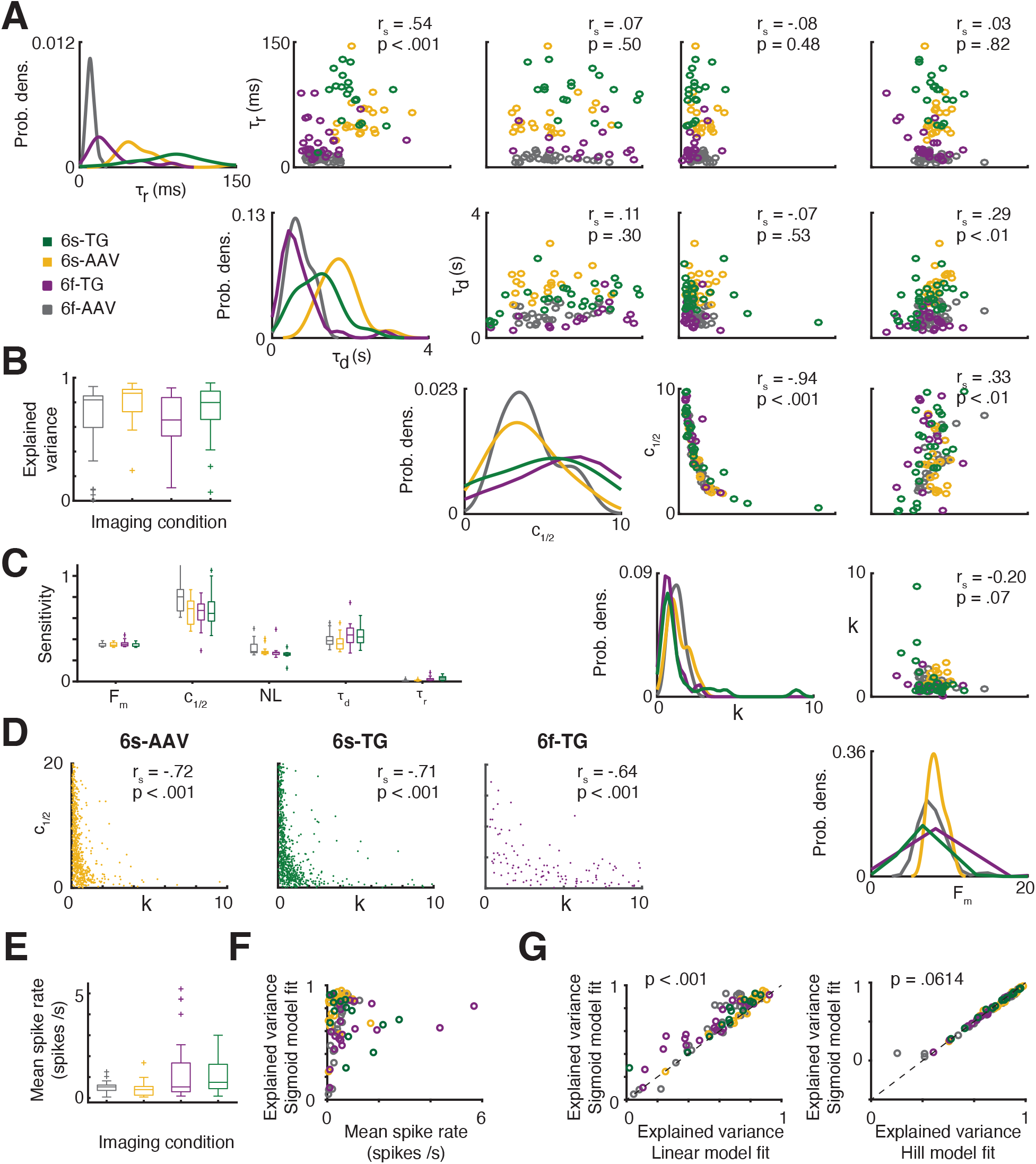
Detailed values of model parameters for simultaneously recorded neurons. **A.** Pairwise correlation plots for each of the spike-to-fluorescence parameters. Panels along the diagonal describe the distribution of each parameter (these are identical to Figure 4B but reproduced to facilitate comparisons). Off-diagonal panels depict the correlation between two parameters. Spearman’s rank correlation of parameters across cells (regardless of recording method) and associated p-value are provided in each off-diagonal panel. Each circle corresponds to a response set. Data from the different indicator conditions is overlaid and marked by color. (gray: 6f-AAV, 11 neurons, 37 response sets; yellow: 6s-AAV, 9 neurons, 21 response sets; purple: 6f-TG, 18 cells, 32 response sets; green: 6s-TG, 22 neurons, 33 recording periods). **B.** Boxplots of explained variance of S2F on validation data for simultaneously recorded neurons (color follows the same convention as in A). **C.** Boxplot of distribution of parameter sensitivity values. **D.** Pairwise correlation of re-estimation of k and c_1/2_ using ALM imaging dynamics (Materials and methods). The re-estimated parameter values are shown as a scatter plot. Each dot corresponds to a neuron (n = 720 for 6s-AAV and 6s-TG; n = 225 for 6f-TG in matched depths). The distribution of the re-estimated parameter values strongly overlapped with those obtained in simultaneous imaging-ephys recordings. c_1/2_ and k had a strong inverse correlation as in the simultaneously recorded data (r_s_ < -.64, p < .001). **E.** Boxplots of firing rates of neurons in each recording sessions (6f-AAV, gray, 0.51 ± 0.25 Hz, mean ± std., range 0.05 – 1.25 Hz; 6s-AAV, yellow, 0.43 ± 0.38 Hz, range 0.05 – 1.68 Hz; 6f-TG, purple, 1.25 ± 1.48 Hz, range 0.09 – 5.22 Hz; 6s-TG, green, 1.08 ± 0.85 Hz, range 0.09 – 3.00 Hz). **F**. Scatter of simultaneous ephys-imaging data model fit and the dynamical range of the data (expressed as mean spike rate). **G.** Scatter of simultaneous ephys-imaging data model fit quality between different S2F models (Materials and methods). Left: comparison between S2F linear model (x-axis) and S2F sigmoid model (y-axis); right, comparison between S2F hill model (x-axis) and S2F sigmoid model (y-axis).

**Figure S4.**
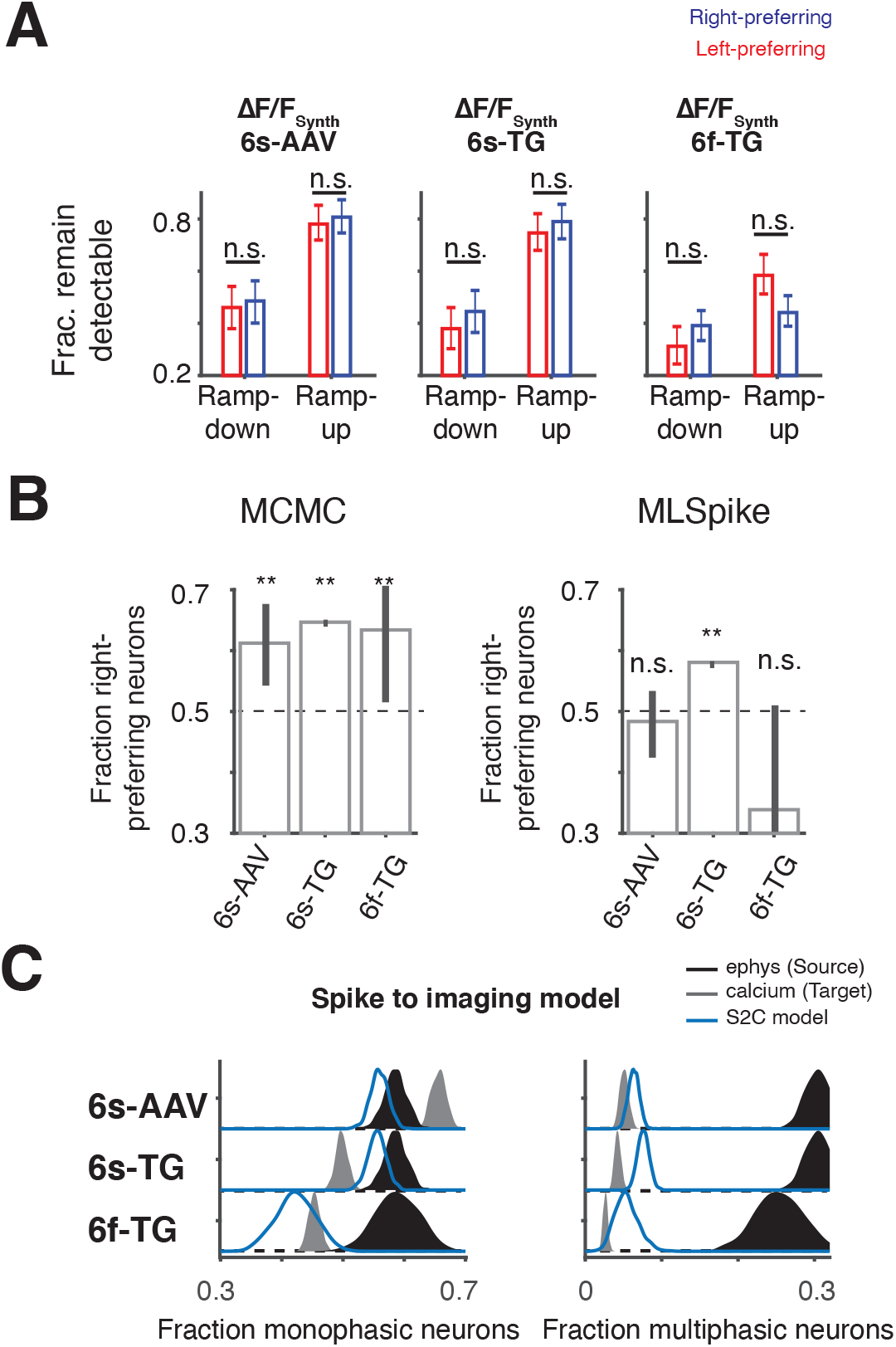
Forward model explains differences in neuronal selectivity between imaging and ephys. **A**. Fraction of cells that remain selective in synthetic imaging plotted separately for ramp-down and ramp-up cells (left: 6s-AAV synthetic, middle: 6s-TG synthetic, right: 6f-TG synthetic), which is further broken down into right- (blue) and left-preferring (red) trials. **B.** Fraction of right-preferring neurons in imaging after spike inference models. Left: the same analyses as that in Figure 2I, but performed on inferred spiking data obtained via the MCMC framework; right: the same analyses as that in Figure 2I, but performed on inferred spiking data obtained via the MLSpike framework. **C.** Estimation of the fraction of monophasic and multiphasic neurons that would be discovered by an imaging experiment through use of the S2F forward model. Plots show the estimates for monophasic (left) and multiphasic (right) neurons. The proportion of the source data, ephys, is in black. The experimentally measured proportions in imaging are in gray. Blue color shows the distribution of selectivity type proportion for different repetitions of each algorithm on subsamples of the dataset for synthetic imaging using 6s-AAV (top), 6s-TG (middle) and 6f-TG (bottom) parameters.

**Figure S5.**
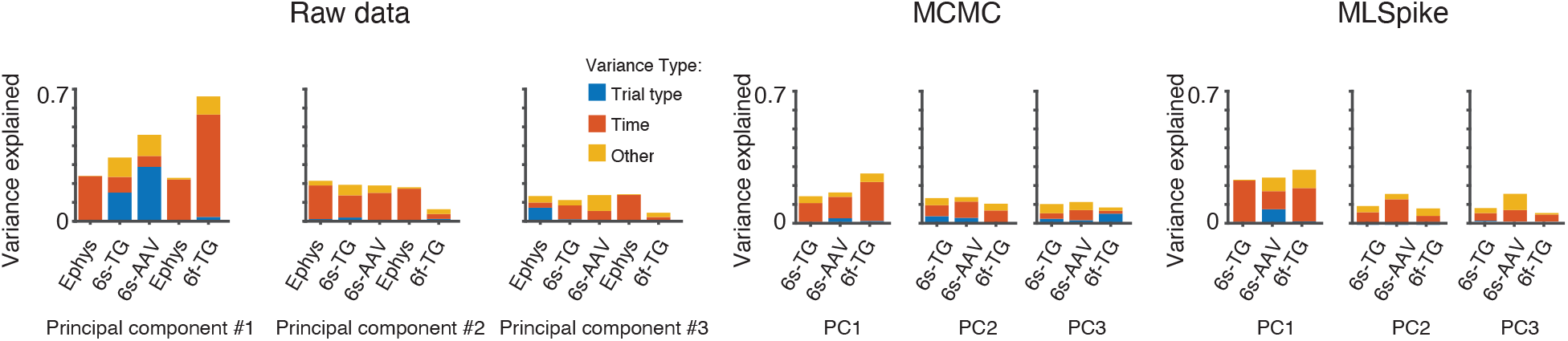
Spike inference results for principal component analysis. Left: fraction of variance explained by principal components 1-3 for each of the datasets, and its division into different sources of variability: red: temporal dynamics; blue: trial type; yellow: other (interaction term). Bars from left to right: ephys, 6s-TG, 6s-AAV; ephys depth-matched to 6f-TG recordings, 6f-TG. Middle: equivalent results for principal component analysis performed on inferred spiking data obtained via the MCMC framework. Right: equivalent results for principal component analysis performed on inferred spiking data obtained via the MLSpike framework.

**Figure S6.**
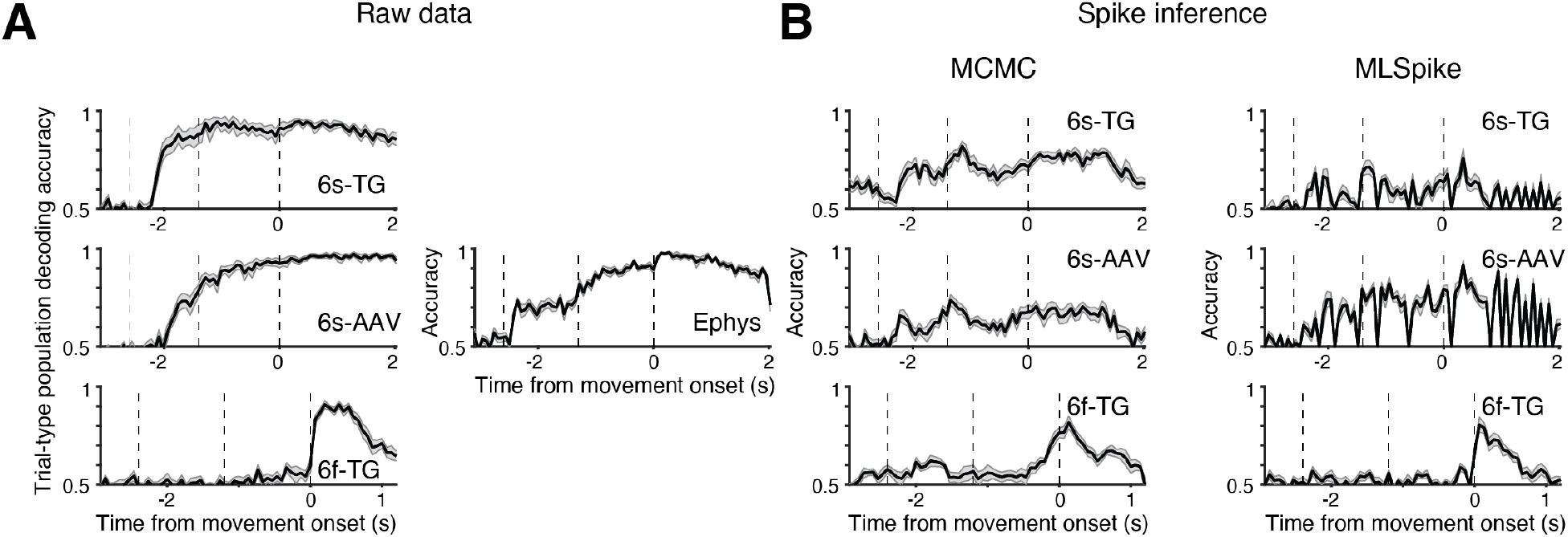
Spike inference results for trial-type population decoding. **A.** Accuracy of trial-type population decoding over time for different datasets. Left column top to bottom: 6s-TG, 6s-AAV, 6f-TG. Right column: ephys. **B.** Accuracy of trial-type population decoding over time of datasets comprised of inferred ephys from the different imaging datasets. top to bottom: 6s-TG, 6s-AAV, 6f-TG. Left column: MCMC framework, right column: MLSpike framework.

**Figure S7.**
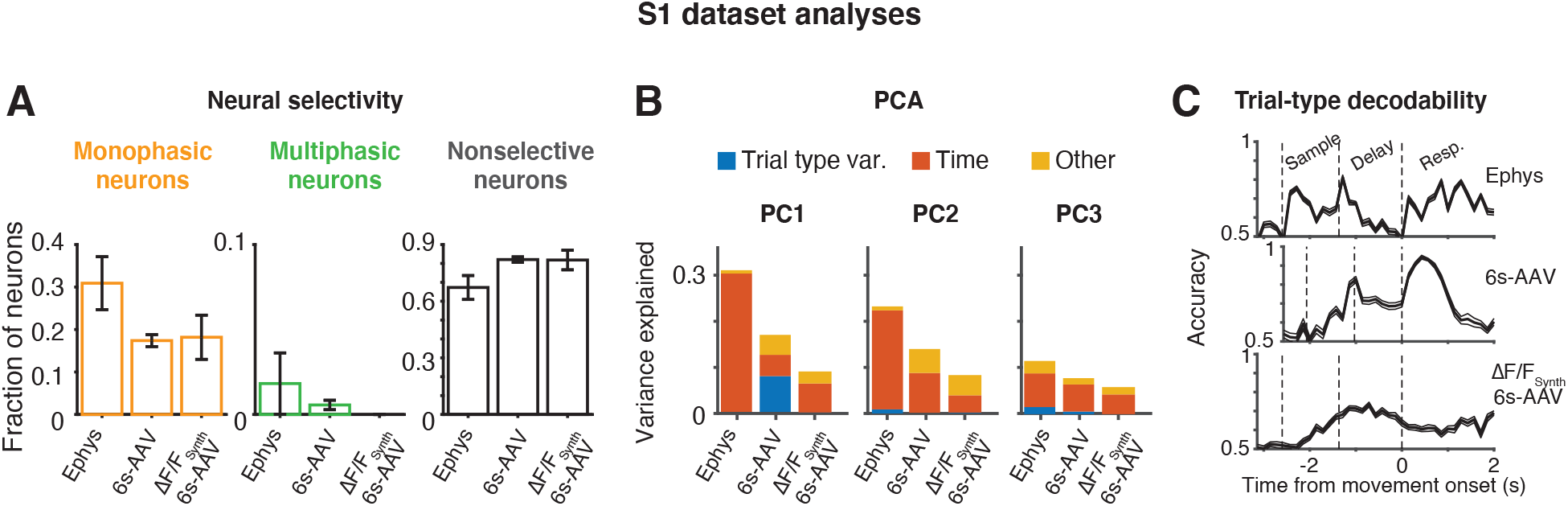
Analysis of matched datasets from a primary somatosensory area. Differences between ephys and imaging are likely to depend not only on the analysis and indicator, but also on the underlying dynamics which change from one brain area to the other. We analyzed a second group of matched population recordings, obtained from primary somatosensory area (S1) rather than ALM. We find that differences in some analyses were no longer present, but others remained. We find that the fraction of multiphasic neurons in S1 was far smaller than that in ALM (n = 1/55, ephys; n = 4/719, 6s-AAV; p < .001, *χ*^2^ test) and there was no significant difference between the fraction of multiphasic neurons observed in ephys and imaging (p = .801, *χ*^2^ test). Our forward model correctly predicted this lack of change (p = .674, *χ*^2^ test between imaging data and synthetic imaging data). Similarly to ALM data, trial type variance dominated the first principal component in imaging but not in ephys and population decoding was substantially delayed in imaging relative to ephys. **A**. Single neuron selectivity type. Bar plots show fraction of neurons found in each of the three selectivity types (left: monophasic, middle: multiphasic, right: nonselective) for the different recording methods (left: ephys, middle: 6s-AAV, right: 6s-AAV synthetic). **B.** principal component variance content. Bar plots show fraction of variance contained in the first three principal components (from left to right: PC1, PC2, PC3). Each bar is broken into the contribution from trial-type variance (blue), time variance (red) and other (yellow). **C.** Population trial-type decodability. Plot shows mean decodability over time for ephys: top, 6s-AAV: middle and synthetic 6s-AAV: bottom. Dashed lines designate different trial periods (sample, delay response). Note that the experiments with 6s-AAV had a slightly shorter delay period, hence the difference in location of dashed lines. Since 6s-AAV synthetic is derived from ephys it has the same trial structure as ephys.

**Figure S8.**
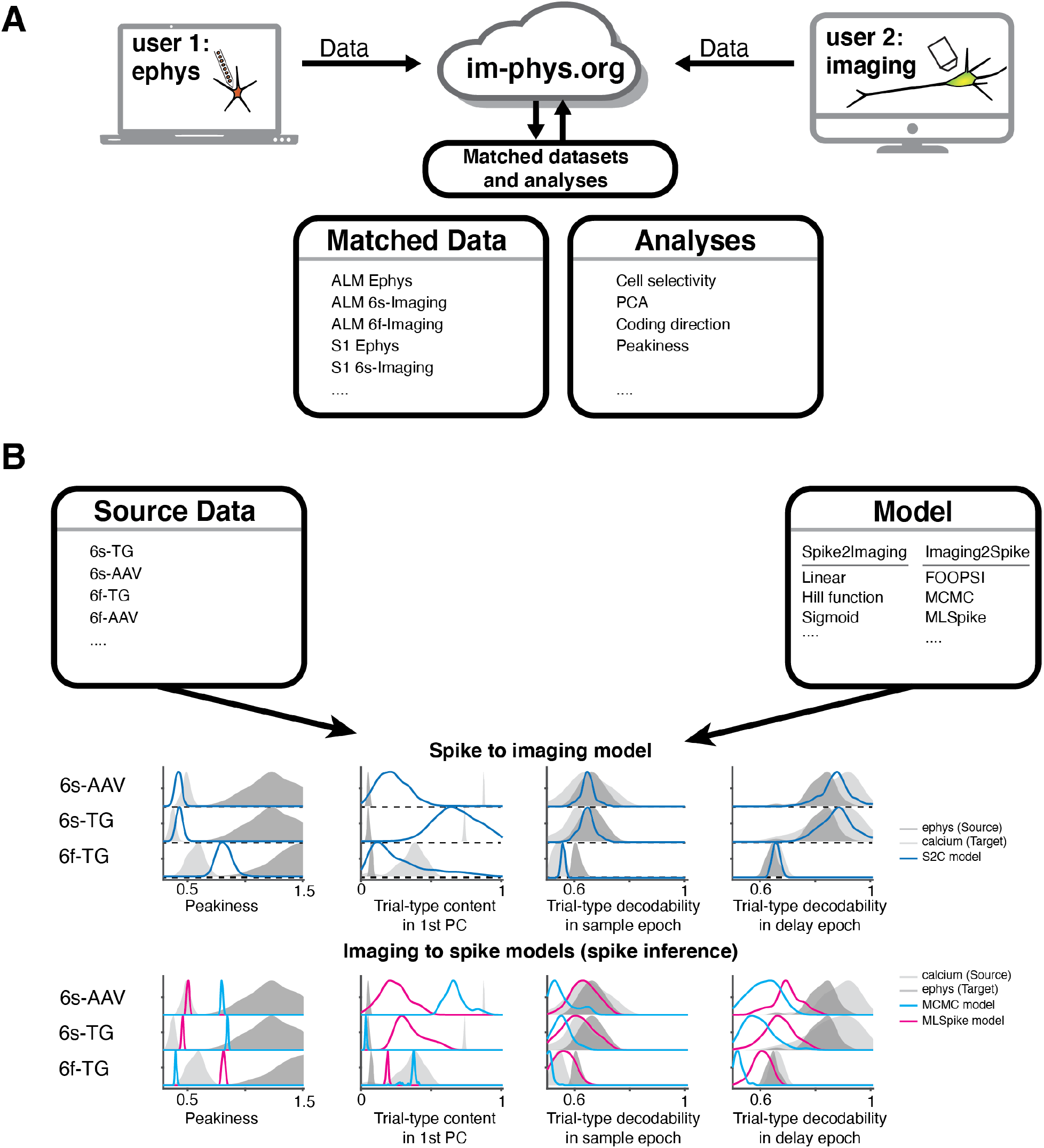
A community based online resource, im-phys.org, for determining quantitative effects of measuring population activity by imaging or ephys. **A.** Top, schematic of our community resource that can allow datasets acquired by different labs to be found in one location and matched in analyses. Bottom, schematic of combining different analyses with different datasets on im-phys.org. **B.** Schematic of using im-phys.org to predict values (metric distributions) expected for different population analyses from datasets acquired by different techniques through use of a variety of forward and inverse models.

**Table S1.**
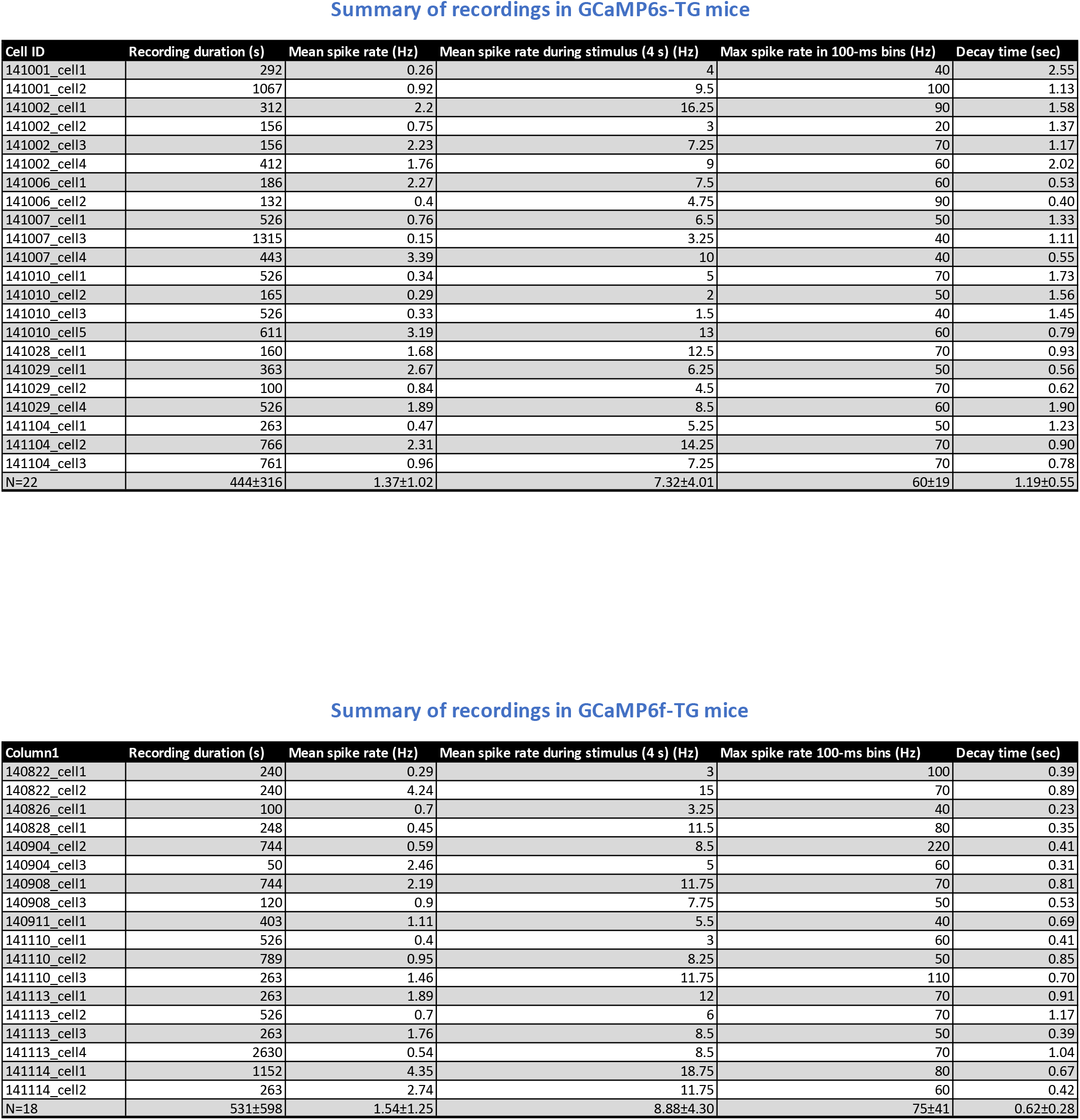

Summary of recordings in GCaMP6s-TG mice
| Cell ID | Recording duration (s) | Mean spike rate (Hz) | Mean spike rate during stimulus (4 s) (Hz) | Max spike rate in 100-ms bins (Hz) | Decay time (sec) |
| --- | --- | --- | --- | --- | --- |
| 141001_cell1 | 292 | 0.26 | 4 | 40 | 2.55 |
| 141001_cell2 | 1067 | 0.92 | 9.5 | 100 | 1.13 |
| 141002_cell1 | 312 | 2.2 | 16.25 | 90 | 1.58 |
| 141002_cell2 | 156 | 0.75 | 3 | 20 | 1.37 |
| 141002_cell3 | 156 | 2.23 | 7.25 | 70 | 1.17 |
| 141002_cell4 | 412 | 1.76 | 9 | 60 | 2.02 |
| 141006_cell1 | 186 | 2.27 | 7.5 | 60 | 0.53 |
| 141006_cell2 | 132 | 0.4 | 4.75 | 90 | 0.40 |
| 141007_cell1 | 526 | 0.76 | 6.5 | 50 | 1.33 |
| 141007_cell3 | 1315 | 0.15 | 3.25 | 40 | 1.11 |
| 141007_cell4 | 443 | 3.39 | 10 | 40 | 0.55 |
| 141010_cell1 | 526 | 0.34 | 5 | 70 | 1.73 |
| 141010_cell2 | 165 | 0.29 | 2 | 50 | 1.56 |
| 141010_cell3 | 526 | 0.33 | 1.5 | 40 | 1.45 |
| 141010_cell5 | 611 | 3.19 | 13 | 60 | 0.79 |
| 141028_cell1 | 160 | 1.68 | 12.5 | 70 | 0.93 |
| 141029_cell1 | 363 | 2.67 | 6.25 | 50 | 0.56 |
| 141029_cell2 | 100 | 0.84 | 4.5 | 70 | 0.62 |
| 141029_cell4 | 526 | 1.89 | 8.5 | 60 | 1.90 |
| 141104_cell1 | 263 | 0.47 | 5.25 | 50 | 1.23 |
| 141104_cell2 | 766 | 2.31 | 14.25 | 70 | 0.90 |
| 141104_cell3 | 761 | 0.96 | 7.25 | 70 | 0.78 |
| N=22 | 444±316 | 1.37±1.02 | 7.32±4.01 | 60±19 | 1.19±0.55 |

Table S1
| Column1 | Recording duration (s) | Mean spike rate (Hz) | Mean spike rate during stimulus (4 s) (Hz) | Max spike rate 100-ms bins (Hz) | Decay time (sec) |
| --- | --- | --- | --- | --- | --- |
| 140822_cell1 | 240 | 0.29 | 3 | 100 | 0.39 |
| 140822_cell2 | 240 | 4.24 | 15 | 70 | 0.89 |
| 140826_cell1 | 100 | 0.7 | 3.25 | 40 | 0.23 |
| 140828_cell1 | 248 | 0.45 | 11.5 | 80 | 0.35 |
| 140904_cell2 | 744 | 0.59 | 8.5 | 220 | 0.41 |
| 140904_cell3 | 50 | 2.46 | 5 | 60 | 0.31 |
| 140908_cell1 | 744 | 2.19 | 11.75 | 70 | 0.81 |
| 140908_cell3 | 120 | 0.9 | 7.75 | 50 | 0.53 |
| 140911_cell1 | 403 | 1.11 | 5.5 | 40 | 0.69 |
| 141110_cell1 | 526 | 0.4 | 3 | 60 | 0.41 |
| 141110_cell2 | 789 | 0.95 | 8.25 | 50 | 0.85 |
| 141110_cell3 | 263 | 1.46 | 11.75 | 110 | 0.70 |
| 141113_cell1 | 263 | 1.89 | 12 | 70 | 0.91 |
| 141113_cell2 | 526 | 0.7 | 6 | 70 | 1.17 |
| 141113_cell3 | 263 | 1.76 | 8.5 | 50 | 0.39 |
| 141113_cell4 | 2630 | 0.54 | 8.5 | 70 | 1.04 |
| 141114_cell1 | 1152 | 4.35 | 18.75 | 80 | 0.67 |
| 141114_cell2 | 263 | 2.74 | 11.75 | 60 | 0.42 |
| N=18 | 531±598 | 1.54±1.25 | 8.88±4.30 | 75±41 | 0.62±0.28 |

